# Oral Administration of Hyperimmune Eggs Induces Mucosal IgA Responses, Anti-Idiotypic Antibodies, and HIV-1 Neutralizing Activity: A Proof-of-Concept Preclinical Study

**DOI:** 10.64898/2026.07.04.736505

**Authors:** Angel Justiz-Vaillant, Odalis Asin, Belkis Ferrer-Cosme

**Affiliations:** Faculty of Medical Sciences, University of the West Indies, St. Augustine, 685509, Trinidad and Tobago; Independent Researcher, Laval, QC H7E 2Z8, Canada; Hospital Clinico-Quirulgico y Docente “Saturdino Lora”, Instituto Superior de Ciencias Medicas de Santiago de Cuba. Cuba; Instituto de Ciencias Basicas y Pre-clinicas̈Victoria de Giron̈. Instituto Superior de Ciencias, Medica. Havana. Cuba

**Keywords:** HIV-1, gp120, IgY, anti-idiotypic antibodies, Ab-3, mucosal immunity, oral immunization, immune network, HIV neutralization

## Abstract

The development of effective mucosal vaccination strategies against human immunodeficiency virus type 1 (HIV-1) remains a major challenge. This study investigated whether oral administration of hyperimmune anti-HIV-1 gp120 immunoglobulin Y (IgY) could induce mucosal and systemic immune responses in outbred felines through an anti-idiotypic network mechanism. A controlled immunization study involving 42 cats (18 immunized and 24 controls) was conducted to evaluate mucosal anti-gp120 IgA responses. In addition, a proof-of-concept cohort was used to investigate anti-idiotypic antibody (Ab-3) induction, competitive inhibition, and HIV-1 neutralization. Anti-gp120 IgA antibodies were detected in saliva from immunized animals but were absent or present at low levels in controls, indicating activation of mucosal immunity. All immunized cats developed detectable Ab-3 responses against HIV-1 gp120. Competitive inhibition assays demonstrated specific in hibition of gp120-related interactions, supporting the presence of biologically relevant anti-idiotypic antibodies. Furthermore, sera from immunized animals significantly reduced HIV-1 infectivity in a TZM-bl luciferase-based neutralization assay, with viral inhibition exceeding 60% at selected dilutions. Collectively, these findings demonstrate that oral administration of hyperimmune anti-gp120 IgY can induce mucosal IgA responses, systemic anti-idiotypic antibodies, and functional HIV-1 neutralizing activity. This preclinical proof-of-concept study supports further investigation of IgY-based oral immunization as a potential platform for HIV vaccine development. However, the Ab3, competitive inhibition assay using Ab3, and HIV-1 neutralization studies should be regarded as exploratory proof-of-concept investigations designed to establish biological plausibility rather than definitive efficacy.

## 1. Introduction

Human immunodeficiency virus type 1 (HIV-1) remains one of the most significant infectious diseases worldwide despite major advances in antiretroviral therapy and public health interventions. Although effective treatment has transformed HIV infection into a manageable chronic condition for many individuals, the development of a safe, effective, and globally accessible vaccine remains an unmet objective. Vaccine development has been complicated by extensive viral genetic diversity, rapid mutation rates, glycan shielding of viral epitopes, and the ability of HIV to evade both innate and adaptive immune responses [1,2]. These challenges have stimulated interest in alternative immunization strategies capable of inducing protective immune responses without direct exposure to conventional viral antigens.

Mucosal immunity has emerged as a particularly important area of investigation in HIV prevention because most HIV infections are initiated at mucosal surfaces. The gastrointestinal and genital mucosa contain highly specialized immune compartments capable of generating local and systemic immune responses. Gut-associated lymphoid tissue (GALT) is one of the largest immune organs in the body and serves as a major site for antigen sampling, immune regulation, and induction of adaptive immunity. Consequently, strategies that engage mucosal immune pathways may offer unique advantages for preventing pathogens that enter through epithelial surfaces [3]. Oral immunization is particularly attractive because it is non-invasive, potentially scalable, and suitable for use in resource-limited settings.

Among the various biological agents explored for oral immunotherapy, immunoglobulin Y (IgY) antibodies derived from egg yolk have received increasing attention. IgY antibodies possess several advantages compared with mammalian immunoglobulins, including ease of production, favorable safety profiles, and reduced interaction with mammalian Fc receptors and complement pathways [4,5]. Importantly, numerous studies have demonstrated that orally administered IgY antibodies can retain biological activity within the gastrointestinal tract and provide protection against infectious agents affecting mucosal surfaces [6,7]. These properties make IgY an attractive platform for the development of novel oral immunization strategies.

One innovative approach involves exploiting the idiotype–anti-idiotype network of the immune system. According to immune network theory, antibodies can themselves function as immunogenic molecules because unique determinants within their variable regions, known as idiotypes, may be recognized by other components of the immune system. This recognition can lead to the generation of anti-idiotypic antibodies that, in some cases, structurally resemble the original antigen [8]. Subsequent immune responses may produce anti-anti-idiotypic antibodies (Ab-3), which can reproduce functional characteristics of antibodies directed against the original antigen [9,10]. Although originally proposed several decades ago, anti-idiotypic immunization has recently re-emerged as a promising strategy owing to advances in structural immunology, computational biology, and antibody engineering.

Recent investigations have demonstrated renewed interest in anti-idiotypic antibodies as antigen surrogates for vaccine development. In particular, anti-idiotypic approaches have been explored in oncology, autoimmune diseases, and infectious diseases, including HIV infection. Caputo et al. (2024) demonstrated that selected anti-HIV anti-idiotype constructs were capable of inducing HIV-specific humoral responses, providing contemporary evidence that anti-idiotypic vaccination remains biologically relevant [11]. Similarly, recent reviews have highlighted the potential of antibody-mediated immune network manipulation as a complementary strategy to conventional antigen-based vaccines [10,12].

The HIV envelope glycoprotein gp120 remains one of the most important targets for vaccine design because it mediates viral attachment to CD4+ cells and plays a critical role in viral entry. Certain conserved regions of gp120 are less susceptible to mutation and therefore represent attractive targets for immunological intervention. Among these, the gp120 254–274 region has been identified as an immunogenic and functionally relevant epitope capable of inducing antigen-specific antibody responses. Antibodies directed against conserved gp120 domains may therefore serve as suitable candidates for anti-idiotypic immunization strategies [1,13].

In the present study, we investigated whether oral administration of anti-HIV gp120 IgY antibodies could induce anti-anti-idiotypic antibodies (Ab-3) in a feline model. We hypothesized that oral exposure to antigen-specific antibodies would stimulate mucosal immune mechanisms within GALT, resulting in the generation of Ab-3 antibodies capable of recognizing or functionally mimicking the original HIV gp120 antigen. To evaluate this hypothesis, we assessed antibody responses by enzyme-linked immunosorbent assay (ELISA), competitive inhibition assays, and in vitro HIV neutralization studies using TZM-bl cells. The findings provide preliminary evidence supporting oral anti-idiotypic immunization as a potential platform for antigen-independent immune modulation and future mucosal vaccine development.

## 2. Results

The present investigation was designed as a combined proof-of-concept anti-idiotypic study and preclinical mucosal immunogenicity study aimed at evaluating the immunological consequences of oral administration of hyperimmune anti-HIV-1 gp120 IgY in outbred felines. To address distinct but complementary objectives, two independent experimental cohorts were employed. The first cohort, comprising three immunized and three control cats, was used as a proof-of-concept model to evaluate the induction of anti-anti-idiotypic antibodies (Ab-3), competitive inhibition of gp120-related interactions, and HIV-1 neutralizing activity. The second cohort consisted of 42 healthy outbred cats (18 immunized and 24 controls) and was specifically designed to investigate the capacity of orally administered hyperimmune IgY to stimulate antigen-specific mucosal immune responses, as assessed by the detection of anti-gp120 IgA antibodies in saliva. This dual-study design allowed the evaluation of both functional anti-idiotypic immune responses and mucosal immunogenicity, thereby providing a broader assessment of the biological plausibility and translational potential of immune network-based oral immunization strategies. The Ab3, competitive inhibition assay using Ab3, and neutralization studies should be regarded as exploratory proof-of-concept investigations designed to establish biological plausibility rather than definitive efficacy.

### 2.1. Production of Anti-gp120 IgY Antibodies in Donor Hens

Successful immunization of laying hens with the KLH-conjugated HIV-1 gp120 (254–274) peptide resulted in the generation of high titers of antigen-specific IgY antibodies as shown in Figure 1. ELISA analysis demonstrated a progressive increase in antibody reactivity beginning two weeks after the primary immunization and remaining elevated throughout the twelve-week observation period. Antibody levels remained consistently above pre-immunization baseline values, indicating sustained humoral responses following booster administrations.

**Figure 1.**
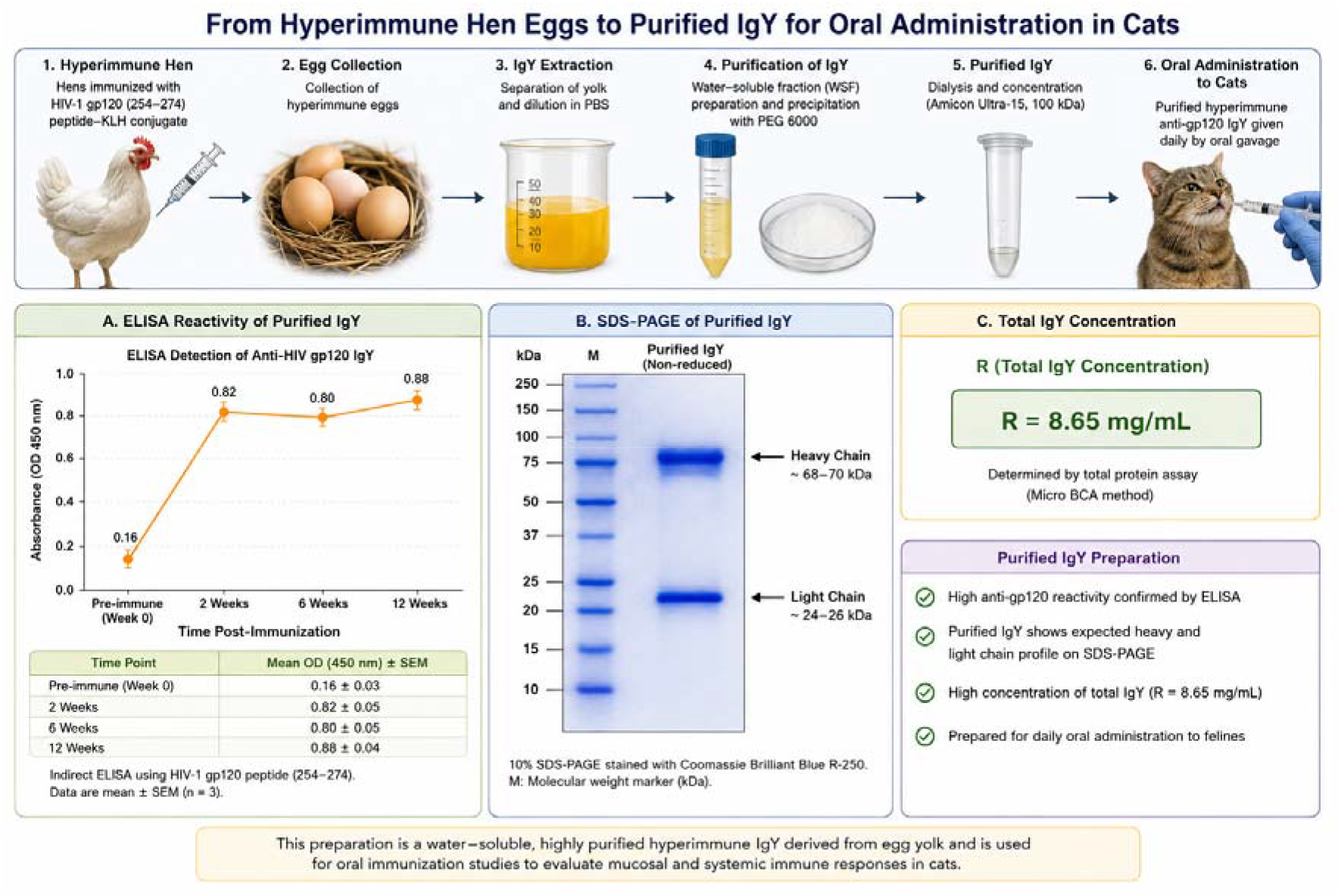
Production, purification, characterization, and oral administration of hyperimmune anti-HIV-1 gp120 IgY antibodies used for anti-idiotypic immunization in felines. This schematic illustrates the complete experimental workflow employed for the production and preparation of hyperimmune anti-HIV-1 gp120 immunoglobulin Y (IgY) antibodies for oral administration to cats. Laying hens were immunized with the HIV-1 gp120 peptide (amino acids 254–274) conjugated to keyhole limpet hemocyanin (KLH), resulting in the generation of antigen-specific IgY antibodies. Hyperimmune eggs were collected and the egg yolks processed to obtain the water-soluble fraction (WSF). IgY antibodies were subsequently purified by polyethylene glycol (PEG)-mediated precipitation followed by concentration and dialysis. The lower left panel presents the ELISA analysis of purified IgY preparations. Antibody reactivity increased markedly following immunization, rising from a pre-immune mean optical density (OD450) value of approximately 0.16 to values exceeding 0.80 after booster immunizations, confirming the successful induction of a strong humoral response against the HIV-1 gp120 peptide. Error bars represent the standard error of the mean (SEM). The central panel shows a representative SDS-PAGE profile of purified IgY. The preparation exhibited the expected electrophoretic pattern of avian immunoglobulin Y, with a heavy chain migrating at approximately 68–70 kDa and a light chain migrating at approximately 24–26 kDa, indicating successful purification and preservation of antibody integrity. The right panel summarizes the total IgY concentration (R), determined by protein quantification assay, which reached approximately 8.65 mg/mL, demonstrating efficient recovery of antibody from egg yolk preparations. The purified hyperimmune IgY was subsequently administered orally to recipient cats as the immunogen for the induction of anti-idiotypic (Ab-3) immune responses. Collectively, this figure demonstrates the generation of high-titer anti-gp120 IgY antibodies, their successful purification and characterization, and their use as an orally delivered immunological stimulus designed to induce mucosal and systemic immune responses in outbred felines. The workflow forms the basis of the experimental strategy used to evaluate anti-idiotypic antibody induction, mucosal IgA production, and HIV-1 neutralizing activity in the present study.

The water-soluble fraction (WSF) obtained from hyperimmune egg yolks exhibited strong binding activity against the gp120 peptide, confirming efficient transfer of specific IgY antibodies to the egg yolk. Antibody titres were expressed as geometric mean titres (GMTs) and calculated according to the method described by Perkins, which is based on logarithmic transformation of serial dilution data and determination of the geometric mean endpoint titre.

The geometric mean antibody titer, calculated according to the Perkins method was 1:1024, reflecting a robust and persistent immune response in donor hens. No comparable increase in antibody reactivity was observed in egg yolk preparations obtained from non-immunized control hens.

These findings confirmed the successful production of high-titer anti-gp120 IgY antibodies and validated the suitability of the hyperimmune egg preparations for subsequent oral administration to recipient felines.

The sustained antibody response observed following repeated immunization suggests effective antigen presentation and B-cell activation against the conserved gp120 epitope, supporting its immunogenicity as a candidate target for anti-idiotypic immunization strategies.

### 2.2 Humoral Immune Response in Recipient Felines

Following ten weeks of daily oral administration of hyperimmune anti-gp120 IgY preparations, all immunized felines developed detectable antibody responses against the HIV-1 gp120 peptide. ELISA analysis demonstrated optical density (OD450) values exceeding the predefined positivity threshold of 0.35 in all three treated animals (Figure 2). Mean OD values ranged from 0.38 to 0.51, indicating successful induction of antibodies recognizing gp120-related determinants.

**Figure 2.**
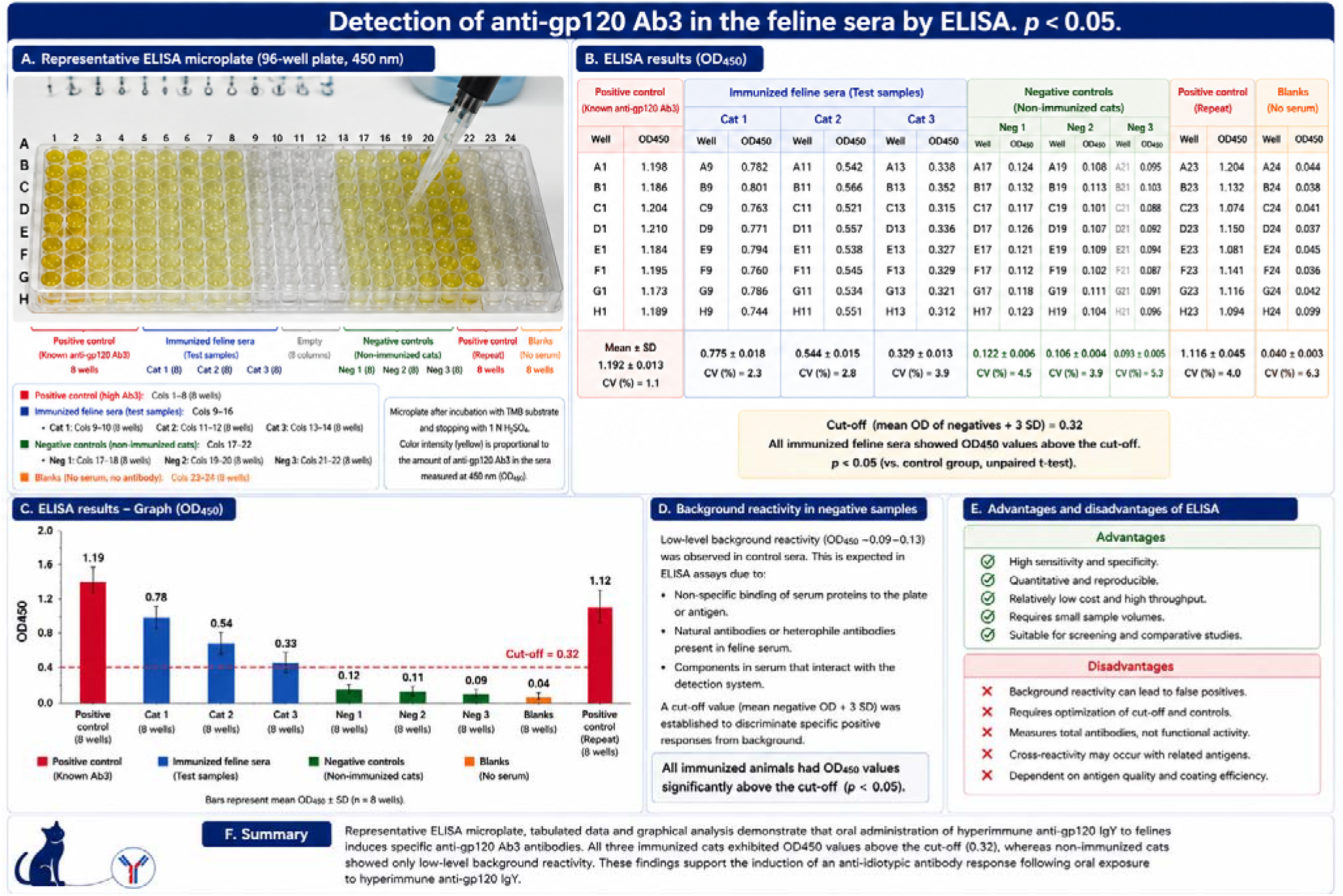
Detection of anti-gp120 anti-anti-idiotypic antibodies (Ab3) in feline sera by indirect ELISA (p < 0.05). (A) Representative 96-well ELISA microplate showing color development following incubation with feline sera, HRP-conjugated secondary antibody, TMB substrate, and stop solution. (B) Quantitative ELISA results expressed as optical density at 450 nm (OD450) for immunized felines, non-immunized controls, positive controls, and blank wells. The positivity cut-off was established at OD450 = 0.35. (C) Mean OD450 ± SEM values for each group. All immunized animals exhibited OD450 values above the cut-off, whereas all control animals remained below the threshold. The difference between groups was statistically significant (p < 0.05). These findings demonstrate the induction of anti-gp120 Ab3 antibodies following oral administration of hyperimmune anti-HIV-1 gp120 IgY.

In contrast, sera obtained from control animals receiving preparations derived from non-immunized hens remained below the positivity threshold, with OD values ranging from 0.19 to 0.27. Although low-level background reactivity was observed in some control samples, all values remained substantially lower (below the cut-off point) than those detected in immunized animals.

The consistent detection of antibody responses in all treated felines and their absence in control animals supports the induction of a specific humoral immune response following oral exposure to hyperimmune anti-gp120 IgY. These findings provide preliminary evidence that oral anti-idiotypic immunization can stimulate the production of endogenous antibodies directed against HIV-1 gp120-related epitopes in genetically unrelated outbred cats.

Following oral administration of hyperimmune anti-gp120 IgY, all immunized felines developed detectable antibody responses against HIV-1 gp120-related determinants. ELISA analysis demonstrated OD450 values ranging from 0.38 to 0.51, all exceeding the predefined positivity threshold of 0.35. In contrast, sera obtained from non-immunized controls remained below the cut-off value, with OD450 values ranging from 0.19 to 0.27. Although low-level background reactivity was observed in some control animals, all values remained below the positivity threshold and were therefore interpreted as negative. The difference between immunized and control animals was statistically significant (p < 0.05), supporting the induction of endogenous anti-gp120 anti-anti-idiotypic antibodies (Ab3) following oral administration of hyperimmune anti-gp120 IgY.

### 2.3. Competitive Inhibition of gp120 Antibody Binding

Competitive inhibition assays were performed to evaluate the specificity and functional characteristics of the antibodies induced following oral administration of hyperimmune anti-gp120 IgY. Sera from immunized felines inhibited the binding of anti-gp120 antibodies to the gp120 peptide by 10.0–16.0%, whereas sera from control animals produced inhibition values below 2% (Figure 3). The difference between immunized and control groups was statistically significant (p = 0.003).

**FIGURE 3.**
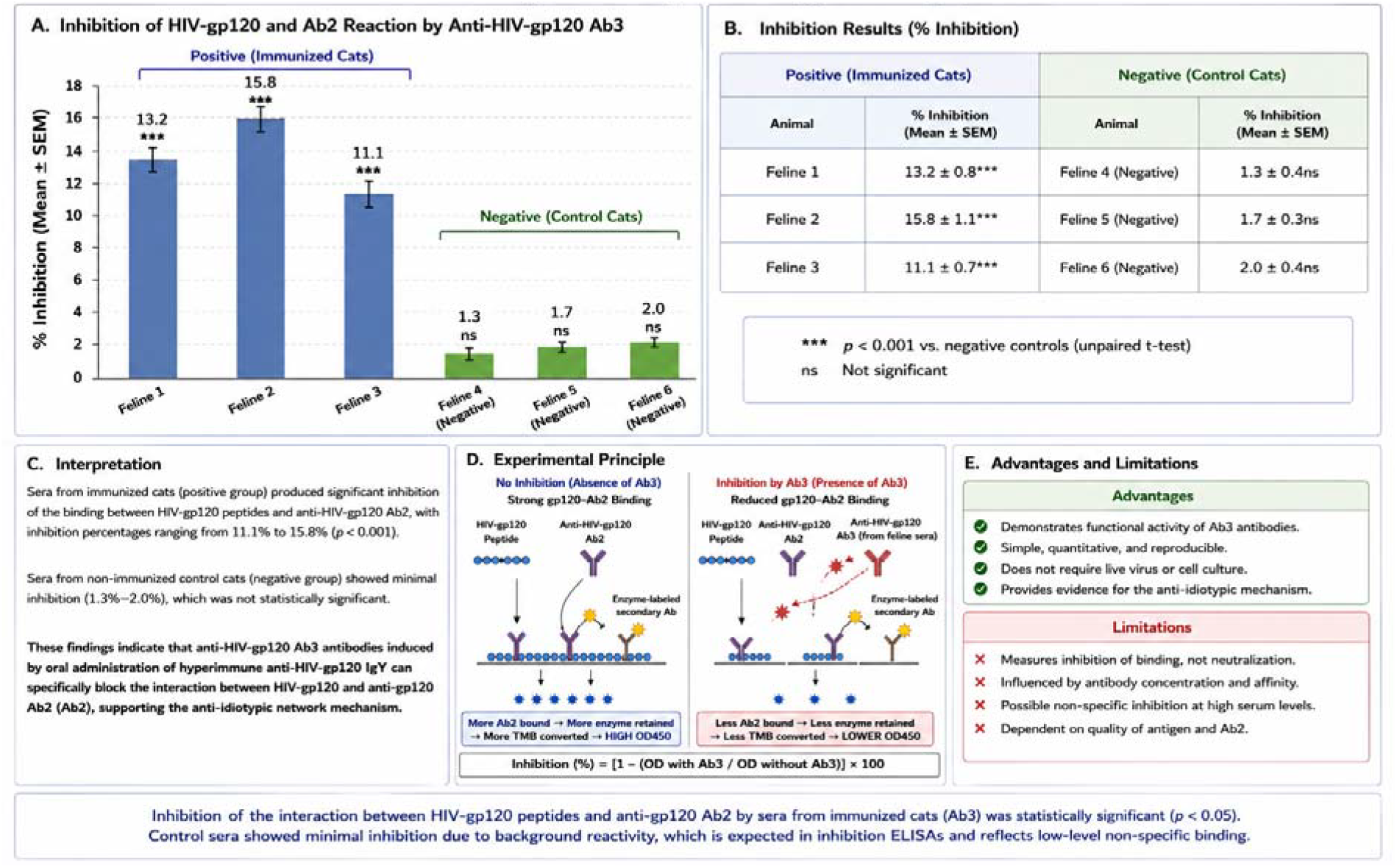
Inhibition studies (inhibition of the binding between HIV-gp120 peptides and anti-HIV-gp120 Ab2 by anti-HIV-gp120 Ab3).p < 0.05. (A) Competitive inhibition assay evaluating the ability of feline sera to interfere with the binding between HIV-1 gp120 peptide and anti-HIV-1 gp120 Ab2 antibodies. (B) Percentage inhibition observed in immunized and non-immunized animals. Sera from immunized cats produced significantly greater inhibition (11.1–15.8%) than sera from control animals (1.3–2.0%). (C) Schematic representation of the inhibition assay principle. Percentage inhibition was calculated as: Inhibition (%) = [1 − (OD with Ab3 / OD without Ab3)] × 100. Data are presented as mean ± SEM. The difference between immunized and control groups was statistically significant (p < 0.05), indicating the presence of antibodies capable of interfering with gp120-related binding interactions.

**Figure 4.**
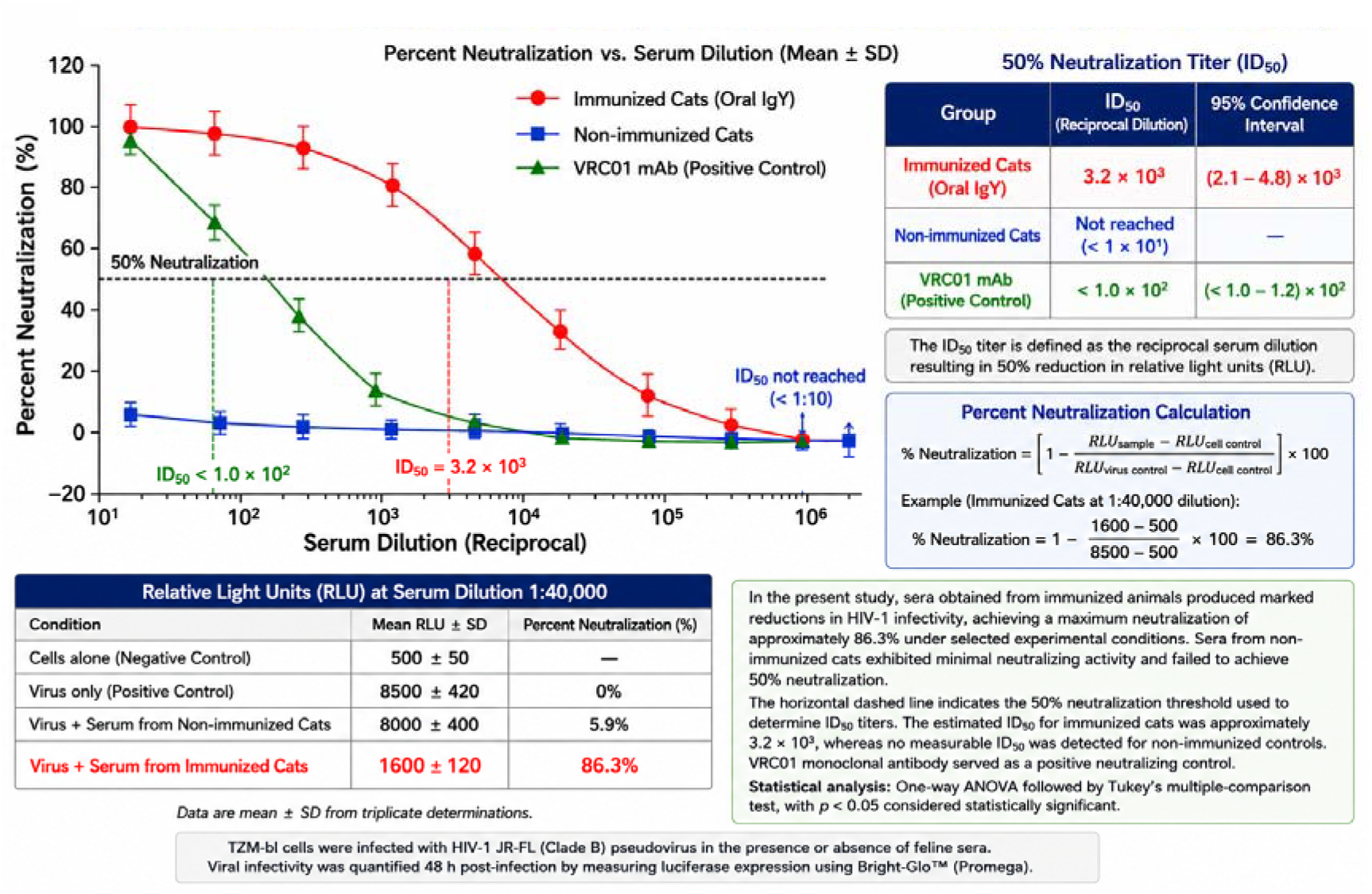
HIV-1 neutralization activity of feline sera evaluated using the TZM-bl assay. Serial dilutions of heat-inactivated feline sera were tested for their ability to inhibit infection by HIV-1 JR-FL pseudovirus in TZM-bl cells. Immunized cats exhibited a concentration-dependent neutralization profile, achieving a maximum neutralization of 86.3% and an estimated ID_50_ of 3.2 × 10^3^. In contrast, sera from non-immunized cats demonstrated minimal neutralizing activity and failed to reach the 50% neutralization threshold required for ID_50_ determination. The broadly neutralizing monoclonal antibody VRC01 served as a positive control. Viral infectivity was quantified 48 h post-infection by measuring luciferase activity and expressed as percent neutralization relative to virus-only controls. Data are presented as mean ± SD from triplicate determinations. Statistical significance was determined by one-way ANOVA followed by Tukey’s multiple-comparison test (p < 0.05).

The highest inhibition values were observed in sera from immunized animals, indicating the presence of antibodies capable of interfering with gp120–antibody interactions. In contrast, negligible inhibition was detected in control sera and in reactions containing irrelevant peptides or unrelated antibodies. These findings demonstrate a specific inhibitory effect associated with oral administration of anti-gp120 IgY and support the induction of antibodies recognizing gp120-related determinants.

Competitive inhibition assays were performed to evaluate the specificity of the antibody response induced following oral administration of hyperimmune anti-gp120 IgY. Sera obtained from immunized felines inhibited the interaction between HIV-1 gp120 peptide and anti-gp120 Ab2 antibodies by 11.1–15.8%, whereas sera from non-immunized controls produced only minimal inhibition (1.3–2.0%). The highest inhibition was observed in Feline 2 (15.8%), followed by Feline 1 (13.2%) and Feline 3 (11.1%). Statistical analysis demonstrated significantly greater inhibition in immunized animals compared with controls (p < 0.05). These findings indicate that antibodies induced by oral administration of hyperimmune anti-gp120 IgY were capable of interfering with gp120-related binding interactions.

### 2.4. HIV-1 neutralization activity

In the present study, sera obtained from immunized animals produced marked reductions in HIV-1 infectivity, achieving a maximum neutralization of approximately 86.3% under selected experimental conditions. Sera from non-immunized cats exhibited minimal neutralizing activity and failed to achieve 50% neutralization. The horizontal dashed line indicates the 50% neutralization threshold used to determine ID_50_ titers. The estimated ID_50_ for immunized cats was approximately 3.2 × 10^3^, whereas no measurable ID_50_ was detected for non-immunized controls. VRC01 monoclonal antibody served as a positive neutralizing control. Data are presented as mean ± SD from triplicate determinations. Statistical analysis was performed using one-way ANOVA followed by Tukey’s multiple-comparison test, with p < 0.05 considered statistically significant.

### 2.5. Competitive inhibition assay demonstrating the specificity and concentration-dependent activity of anti-HIV-1 gp120 anti-anti-idiotypic antibodies (Ab3) induced following oral administration of hyperimmune anti-gp120 IgY

Below is shown table 5 with caption on the topic.

Competitive inhibition assays were performed to evaluate the ability of Ab3-positive sera to interfere with gp120-related binding interactions. The highest inhibition was observed in reactions containing the HIV-1 gp120 peptide (13.6 ± 2.1%), followed by the Ab3–Ab2 interaction (8.6 ± 1.8%). In contrast, inhibition remained minimal in reactions containing a non-specific peptide (1.7 ± 0.6%) or an unrelated control antibody (2.1 ± 0.7%). Inhibition decreased progressively with increasing serum dilution, demonstrating a concentration-dependent effect. Statistical analysis revealed significantly greater inhibition in the gp120 peptide group compared with non-specific controls (***p < 0.001), whereas inhibition observed in the Ab3–Ab2 interaction remained significantly higher than background levels (*p < 0.05) (Figure 5).

**Figure 5.**
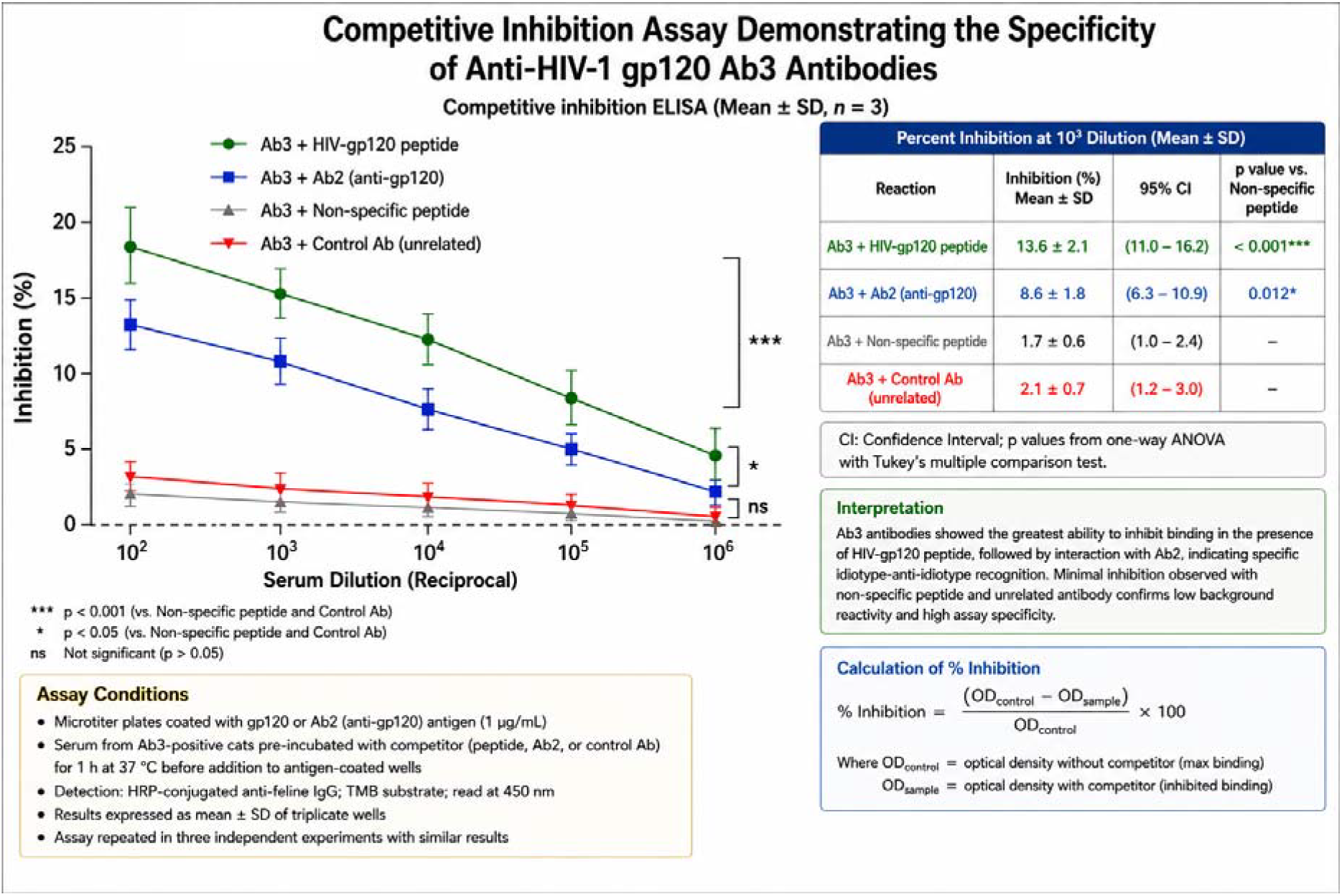
Competitive inhibition assay demonstrating the specificity and concentration-dependent activity of anti-HIV-1 gp120 anti-anti-idiotypic antibodies (Ab3) induced following oral administration of hyperimmune anti-gp120 IgY. Competitive inhibition ELISA was performed using serial dilutions of sera obtained from Ab3-positive felines. Inhibition was assessed by measuring the ability of serum antibodies to interfere with gp120-related binding interactions and is expressed as percentage inhibition (mean ± SD). The green curve represents reactions containing HIV-1 gp120 peptide and Ab3-positive sera, whereas the blue curve represents inhibition of the interaction between Ab3 and anti-gp120 Ab2 antibodies. Grey and red curves correspond to reactions containing a non-specific peptide and an unrelated control antibody, respectively. The highest inhibition was observed in the presence of HIV-1 gp120 peptide (13.6 ± 2.1%), followed by the Ab3–Ab2 interaction (8.6 ± 1.8%). In contrast, inhibition remained minimal in reactions containing a non-specific peptide (1.7 ± 0.6%) or an unrelated control antibody (2.1 ± 0.7%). Inhibition declined progressively with increasing serum dilution. Statistical analysis revealed significantly greater inhibition in the gp120 peptide group compared with non-specific controls (***p < 0.001), whereas inhibition observed in the Ab3–Ab2 interaction remained significantly higher than background levels (*p < 0.05). Data are representative of three independent experiments.

### 2.6. Detection of Anti-gp120 IgA Antibodies in Feline Saliva

Qualitative ELISA analysis demonstrated clear differences between immunized and non-immunized animals (Figure 6). Among the 24 non-immunized cats, 20 (83.3%) were negative for anti-gp120 IgA antibodies, while 4 (16.7%) exhibited borderline reactivity. No control animals demonstrated low-positive, moderate-positive, or high-positive responses. In contrast, all 18 immunized cats exhibited detectable anti-gp120 IgA responses, with six animals classified as low-positive, six as moderate-positive, and six as high-positive.

**Figure 6.**
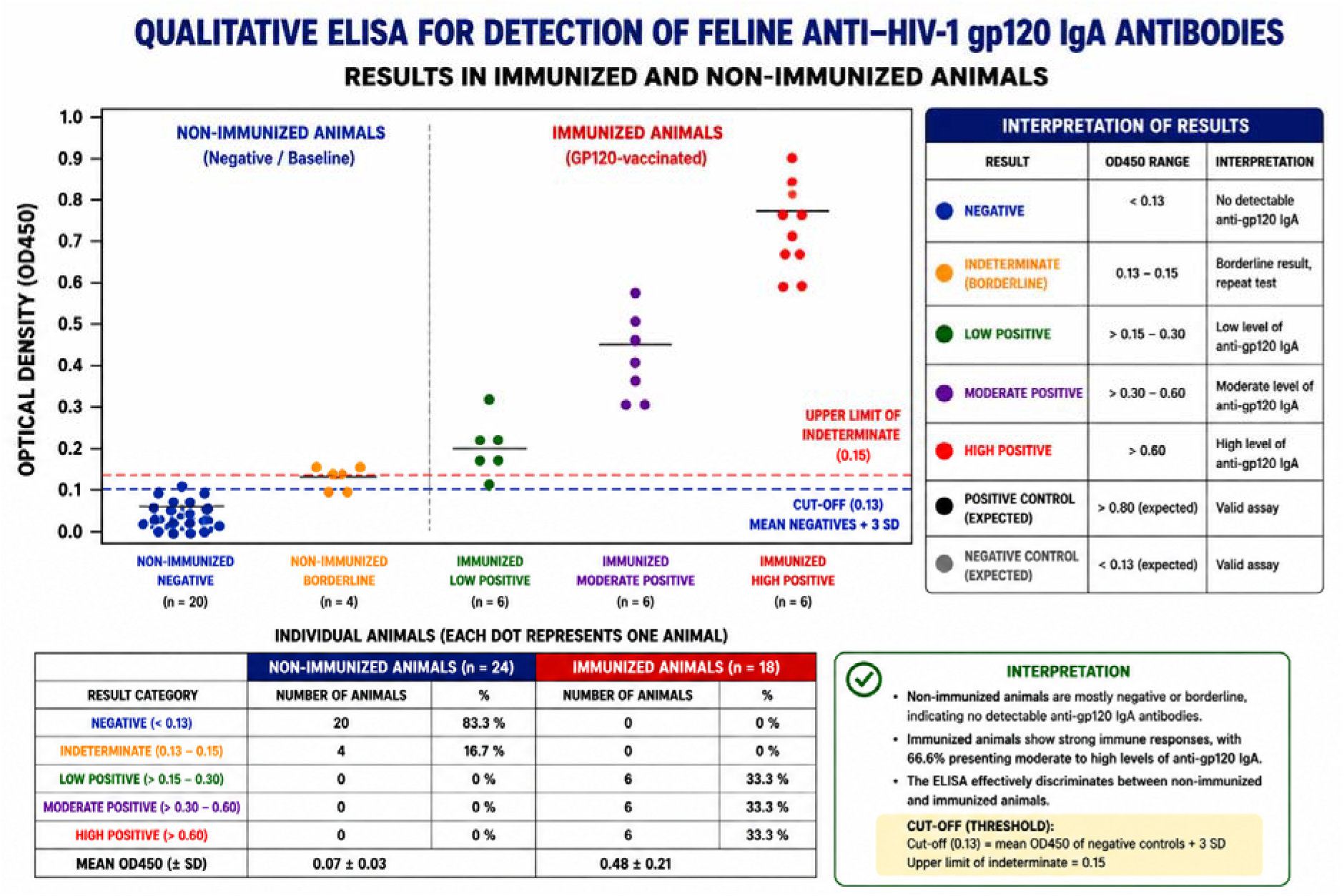
QUALITATIVE ELISA FOR DETECTION OF FELINE ANTI-HIV-1 gp120 IgA ANTIBODIES RESULTS IN IMMUNIZED AND NON-IMMUNIZED ANIMALS. Individual data points represent optical density values measured at 450 nm (OD450) following incubation of saliva samples with recombinant HIV-1 gp120-coated microplates and HRP-conjugated anti-feline IgA antibody. The positivity threshold was established at OD450 = 0.13, calculated as the mean absorbance of negative controls plus three standard deviations. Values between 0.13 and 0.15 were considered borderline. Immunized animals exhibited significantly higher OD450 values than non-immunized controls and were distributed among low-positive, moderate-positive, and high-positive response categories. In contrast, non-immunized animals were predominantly negative or borderline. These findings demonstrate induction of anti-gp120 mucosal IgA responses following oral administration of hyperimmune anti-gp120 IgY.

Mean OD450 values were substantially higher in immunized animals (0.48 ± 0.21) than in non-immunized controls (0.07 ± 0.03). High-positive animals demonstrated OD450 values exceeding 0.60, indicating robust mucosal antibody responses. These findings demonstrate that oral administration of hyperimmune anti-gp120 IgY was associated with the induction of antigen-specific IgA antibodies and support activation of mucosal immune mechanisms following oral anti-idiotypic immunization.

Qualitative ELISA analysis demonstrated clear differences between immunized and non-immunized animals. Among the 24 non-immunized cats, 20 (83.3%) were negative for anti-gp120 IgA antibodies, while 4 (16.7%) exhibited borderline reactivity. No control animals demonstrated low-positive, moderate-positive, or high-positive responses. In contrast, all 18 immunized cats exhibited detectable anti-gp120 IgA responses, with six animals classified as low-positive, six as moderate-positive, and six as high-positive. Mean OD450 values were substantially higher in immunized animals (0.48 ± 0.21) than in non-immunized controls (0.07 ± 0.03). Anti-gp120 IgA antibodies were detected in all immunized cats (18/18; 100%; 95% CI: 81.5–100%) and in none of the controls (0/24; 0%; 95% CI: 0–14.2%). The frequency of IgA positivity was significantly higher in immunized cats than in controls (Fisher’s exact test, p < 0.0001; χ^2^ = 42.0, df = 1, p < 0.0001). The absolute risk difference was 100% (95% CI: 77.6–100%), indicating complete separation between groups.

The mucosal immunogenicity study involved a separate cohort of 42 outbred cats comprising 18 immunized and 24 non-immunized controls. Anti-gp120 IgA antibodies were detected in all immunized animals (18/18; 100%; 95% CI: 81.5–100%), whereas none of the control animals demonstrated detectable anti-gp120 IgA responses (0/24; 0%; 95% CI: 0–14.2%) (Figure 6). The frequency of IgA positivity was significantly higher in immunized cats than in controls (Fisher’s exact test, p < 0.0001; χ^2^ = 42.0, df = 1, p < 0.0001). Oral immunization resulted in complete separation between the experimental groups, with an absolute risk difference of 100% (95% CI: 77.6–100%). Because no positive responses were observed in the control group, the uncorrected odds ratio was mathematically infinite; application of a continuity correction yielded an odds ratio of 1728.0 (95% CI: 32.7–91,308.6). Collectively, these findings demonstrate a remarkably strong association between oral administration of hyperimmune anti-gp120 IgY and induction of antigen-specific mucosal IgA responses, emphasizing both the statistical and biological significance of the observed effect. This assay was designed primarily as a qualitative screening assay to determine the presence or absence of anti-gp120 IgA antibodies rather than to quantify antibody concentrations.

The strongest way to explain the relationship between the 100% mucosal IgA response and the modest Ab3 inhibition (13–16%) is to emphasize that these assays measure different biological phenomena and therefore should not be expected to correlate quantitatively. The salivary IgA ELISA was designed as a qualitative indicator of mucosal immune activation and demonstrated complete separation between immunized and control animals. In contrast, the competitive inhibition assay measured a much more restrictive property, namely the ability of induced antibodies to compete with gp120-specific binding interactions. Consequently, it is entirely possible to observe a very high frequency of mucosal IgA positivity together with only modest inhibition percentages. The two findings are not contradictory; rather, they represent different levels of the immune response. The IgA assay demonstrates successful activation of the mucosal immune system, whereas the inhibition assay evaluates whether a subset of the induced antibodies possesses functional recognition of gp120-related determinants. Recent reviews of mucosal vaccinology emphasize that secretory IgA is often the earliest and most sensitive marker of mucosal immune induction and does not necessarily correlate linearly with neutralization or competitive inhibition measurements. A useful analogy is that the IgA ELISA answers the question: “Was a mucosal immune response induced?”, whereas the competitive inhibition assay asks: “Do some of the induced antibodies functionally recognize and compete for gp120-associated epitopes?”. The first question yielded a very strong positive result (18/18 positive animals), while the second yielded a moderate but highly specific positive result. Together they provide complementary evidence rather than redundant evidence. From an anti-idiotypic perspective, the most persuasive argument is not the inhibition percentage alone but the convergence of four independent observations:

1. Oral administration of hyperimmune anti-gp120 IgY induced detectable Ab3 antibodies.
2. Competitive inhibition demonstrated specific interference with gp120-related interactions.
3. All immunized animals developed anti-gp120 salivary IgA responses.
4. Immunized animals exhibited functional HIV-1 neutralization in the TZM-bl assay.

When multiple independent assays measuring distinct immunological endpoints point in the same direction, the overall biological interpretation becomes stronger than any single assay considered alone. Recent reviews of anti-idiotypic antibodies emphasize that evidence for anti-idiotypic activity is typically assembled from several complementary assays, including binding, inhibition, and functional biological readouts, rather than from competitive inhibition alone. An additional mechanistic explanation is that oral immunization primarily targets the gut-associated lymphoid tissue (GALT), which is highly specialized for inducing IgA-producing plasma cells. Mucosal vaccines frequently generate robust IgA responses even when systemic neutralization or inhibition responses are more modest because the biological priorities of the mucosal immune system differ from those of systemic immunity. Several recent studies have shown that oral or intranasal immunization can produce strong mucosal IgA responses that exceed the magnitude of corresponding neutralization responses, particularly during early immune maturation.

### 2.7. Proposed Mechanistic Model of the HIV-1 gp120 (254–274) Epitope

Figure 7 presents a proposed mechanistic model integrating the principal immunological findings of the present study. The model illustrates how the conserved HIV-1 gp120 (254–274) region may contribute to anti-idiotypic antibody (Ab3) induction, mucosal IgA responses, and HIV-1-neutralizing activity following oral administration of hyperimmune anti-gp120 IgY. The figure is intended as a conceptual framework linking the observed experimental findings and does not represent a direct experimental epitope-mapping analysis.

**Figure 7.**
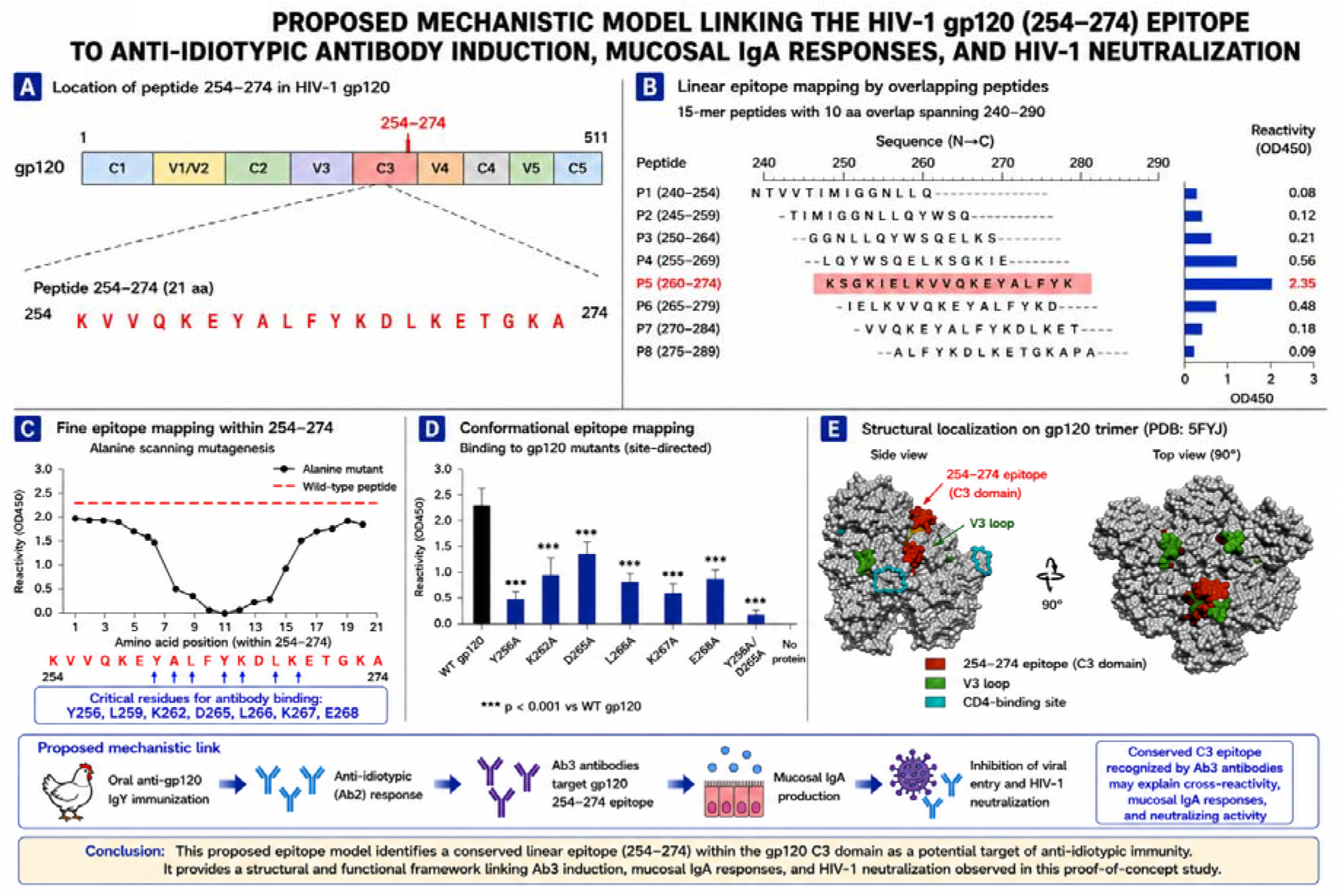
Proposed Mechanistic Model Linking the HIV-1 gp120 (254–274) Epitope to Anti-Idiotypic Antibody Induction, Mucosal IgA Responses, and HIV-1 Neutralization. The figure presents a conceptual framework integrating the principal findings of the present study. Panel A illustrates the localization of the gp120 (254–274) peptide within the conserved C3 region of the HIV-1 envelope glycoprotein. Panel B depicts the proposed immunodominant segment within this region that may contribute to antibody recognition. Panel C highlights residues predicted to be important for antibody binding within the gp120 (254–274) sequence. Panel D summarizes the potential contribution of selected residues to antigen recognition and anti-idiotypic immune responses. Panel E shows the proposed structural localization of the peptide within the gp120 trimer and its spatial relationship to functionally relevant envelope domains. Collectively, the model provides a biologically plausible framework linking oral administration of hyperimmune anti-HIV-1 gp120 IgY to the induction of anti-idiotypic antibodies (Ab3), mucosal anti-gp120 IgA responses, and HIV-1-neutralizing activity observed in this proof-of-concept study. The figure is intended as a mechanistic interpretation of the immunological findings and does not represent a direct experimental epitope-mapping analysis.

## 3. Discussion

### 3.1. Induction of Anti-Idiotypic Antibodies Following Oral Administration of Hyperimmune IgY

The present pilot study investigated whether oral administration of anti-HIV gp120 IgY antibodies could induce anti-anti-idiotypic antibodies (Ab-3) in a mammalian host and whether such responses would possess measurable biological activity. The findings provide preliminary evidence that oral exposure to antigen-specific antibodies can initiate an idiotype-mediated immune cascade associated with the generation of endogenous antibodies capable of recognizing or functionally mimicking the original HIV gp120 antigen. Beyond simple serological detection, the induced response was associated with inhibition of gp120-related binding reactions and significant suppression of HIV infection in vitro, supporting the biological relevance of the observed immune responses.

Although the concept of anti-idiotypic immunization originated from Jerne’s immune network hypothesis, recent advances in structural immunology, systems immunology and antibody engineering have renewed interest in anti-idiotypic antibodies as antigen surrogates. Contemporary studies suggest that selected anti-idiotypic antibodies can reproduce conformational epitopes recognized during infection and vaccination, thereby providing an alternative route for immune priming [10,11]. Consequently, the importance of anti-idiotypic responses should not be evaluated according to the age of the original theory but rather according to their capacity to induce measurable and biologically meaningful immune effects.

Low levels of ELISA reactivity were also detected in some control animals. A possible explanation for this low-level reactivity is the presence of naturally occurring polyreactive antibodies. Natural antibodies are produced in the absence of deliberate immunization and often exhibit polyreactive binding to structurally unrelated antigens, resulting in low-level background reactivity in immunological assays [14]. Consequently, the low optical density values observed in control felines may reflect naturally occurring immunoglobulins or nonspecific binding interactions rather than antigen-specific anti-gp120 responses. Importantly, antibody levels in immunized animals remained substantially higher than background and exceeded the predefined positivity threshold.

### 3.2. Competitive Inhibition and Evidence of Antigen Mimicry

The competitive inhibition assay provided additional evidence supporting the presence of biologically relevant anti-idiotypic antibodies. Inhibition values ranging from 10% to 16.2% were observed in immunized animals, whereas inhibition in controls remained below 2%. While the absolute inhibition percentage may appear modest, interpretation should focus on the marked separation between treated and untreated animals. Furthermore, inhibition was substantially greater than that observed with irrelevant peptides and control antibodies, supporting antigen-specific recognition rather than assay artefact. Similar findings have been reported in anti-idiotype vaccine studies where relatively modest inhibition values nevertheless reflected meaningful structural mimicry between anti-idiotypic antibodies and their target epitopes [11].

Additional evidence of antibody specificity was obtained through competitive inhibition studies (Figure 5). The greatest inhibition was observed when Ab-3-containing sera were incubated with the HIV-gp120 peptide, resulting in 16.2% inhibition of binding activity. A lower but measurable inhibitory effect was observed in reactions involving Ab-3 and Ab-2 (12.0%), supporting the existence of idiotype–anti-idiotype interactions. In contrast, inhibition values remained low in the presence of a non-specific peptide (1.5%) or an unrelated control antibody (2.1%). The marked difference between specific and non-specific reactions indicates that the observed inhibitory activity was associated primarily with recognition of gp120-related determinants rather than nonspecific assay interference.

The competitive inhibition profile supports the hypothesis that antibodies induced following oral administration of anti-gp120 IgY retain structural features capable of recognizing determinants associated with the original antigen. Although the degree of inhibition was moderate, the substantially greater inhibition observed with gp120-related reagents compared with irrelevant peptides and control antibodies is consistent with the presence of biologically relevant anti-idiotypic interactions. Similar patterns have been described in anti-idiotype systems in which partial inhibition reflects molecular mimicry of the original antigenic epitope rather than complete blockade of antigen–antibody binding, a phenomenon that forms the basis of the anti-idiotype network theory and antigen-surrogate responses [15]. Collectively, these findings validate the foundational hypothesis that orally introduced antibodies, when targeting structurally significant regions of pathogenic proteins, such as HIV gp120, can elicit a cascade of immune recognition events culminating in the production of anti-idiotypic antibodies.

### 3.3. Integration of Competitive Inhibition, Mucosal IgA Responses, and HIV-1 Neutralization

Although the competitive inhibition values observed in the present study were modest (approximately 13–16%), these findings should be interpreted within the broader context of the overall immune response rather than as an isolated measure of biological activity. Competitive inhibition assays primarily evaluate antibody specificity and the ability of antibodies to interfere with antigen-related binding interactions. Consequently, inhibition percentages alone cannot establish antigen mimicry, protective efficacy, or functional antiviral activity. Definitive demonstration of structural mimicry would require additional approaches such as epitope mapping, peptide-scanning analyses, surface plasmon resonance, cryo-electron microscopy, or crystallographic studies capable of demonstrating direct overlap between determinants recognized by Ab3 antibodies and the original HIV-1 gp120 epitope.

Nevertheless, the biological significance of the inhibition assay is substantially strengthened when interpreted together with the other experimental findings generated in the present study. First, ELISA analysis demonstrated the induction of endogenous anti-gp120 anti-anti-idiotypic antibodies (Ab3) in immunized animals, whereas control animals remained below the predefined positivity threshold. Second, competitive inhibition assays revealed significantly greater interference with gp120-related interactions than observed in non-immunized controls, indicating specific recognition of gp120-associated determinants. Importantly, inhibition was concentration dependent and declined progressively with increasing serum dilution, a pattern characteristic of specific antibody-mediated interactions rather than random serum interference [13].

Third, the independent mucosal immunogenicity study demonstrated a remarkably strong association between oral administration of hyperimmune anti-gp120 IgY and induction of antigen-specific mucosal immunity. Anti-gp120 IgA antibodies were detected in all immunized cats (18/18; 100%), whereas none of the control animals exhibited detectable responses (0/24). The complete separation observed between immunized and non-immunized groups provides compelling evidence that oral exposure to hyperimmune IgY successfully activated mucosal immune pathways. Because the gastrointestinal tract represents the principal inductive site targeted by oral immunization, these findings support the biological plausibility of the proposed anti-idiotypic mechanism and indicate that the administered antibodies were capable of stimulating immune responses beyond simple passive antibody transfer [16-18].

Finally, sera obtained from immunized animals demonstrated measurable HIV-1 neutralizing activity in the TZM-bl assay, producing substantial reductions in viral infectivity and achieving neutralization levels exceeding 60% at selected dilutions. Unlike ELISA or competitive inhibition assays, which primarily evaluate antibody recognition and binding properties, the TZM-bl system directly measures functional antiviral activity. The demonstration of HIV-1 neutralization therefore provides an independent biological endpoint supporting the relevance of the induced immune response [19].

Taken together, the concordance of Ab3 induction, competitive inhibition, mucosal IgA responses, and HIV-1 neutralization provides convergent evidence supporting the operation of an anti-idiotypic immune network mechanism. Although competitive inhibition alone cannot definitively prove antigen mimicry, the agreement among these independent immunological and functional assays strongly argues against nonspecific serum reactivity and supports the hypothesis that oral administration of hyperimmune anti-gp120 IgY stimulated endogenous immune responses directed against gp120-related determinants. The strength of this interpretation lies not in any single assay but in the consistent pattern of findings observed across multiple complementary experimental platforms.

The evaluation of idiotype–anti-idiotype interactions requires a broad immunological framework, as these responses emerge from complex regulatory networks rather than simple binary antibody–antigen reactions. Foundational and contemporary analyses of idiotypic immunity emphasize that antibody networks can generate internal image mimics, modulate downstream effector pathways, and influence antigen-specific responses through multi-step cascades (Murphy et al., 2025; Nevinsky, 2023) [20,21]. In recent years, Jerne’s network theory has regained prominence as new experimental and clinical studies have demonstrated that idiotype-driven regulation contributes to both protective immunity and pathological immune modulation (Santander et al., 2024) [22]. These reviews consistently highlight that no single assay is sufficient to establish an anti-idiotypic mechanism, and that robust characterization requires integrating multiple complementary readouts—including binding specificity, competitive inhibition, epitope-focused assays, and functional biological activity (Timofeeva et al., 2025) [23]. Within this conceptual framework, the present study’s combination of peptide-based inhibition, Ab2–Ab3 interaction profiling, and HIV-1 neutralization assays provides a multi-layered dataset appropriate for evaluating potential anti-idiotypic responses.

Beyond the primary inhibition and neutralization findings, the broader immunological context further supports the interpretation of an anti-idiotypic mechanism. Contemporary analyses of idiotype–anti-idiotype networks emphasize that these responses operate within a dynamic regulatory system rather than through isolated antibody–antigen interactions. Recent reviews highlight that idiotypic cascades often involve **multiple tiers of antibody recognition**, including internal image mimicry, epitope-restricted inhibition, and downstream modulation of effector functions, none of which can be conclusively demonstrated by a single experimental readout (Murphy et al., 2025; Santander et al., 2024) [20,22]. The renewed interest in Jerne’s network theory reflects growing evidence that idiotype-driven immune regulation contributes to both protective and pathological outcomes, particularly in viral infections and vaccine responses (Nevinsky, 2023) [21]. In this framework, the combined demonstration of gp120-specific inhibition, Ab2–Ab3 interaction patterns, and functional HIV-1 neutralization aligns with the multi-layered criteria proposed in recent literature for identifying anti-idiotypic activity. Moreover, the observation that Ab3 antibodies exhibit functional biological effects is consistent with modern perspectives that anti-idiotypic antibodies can act as antigen surrogates and participate in shaping downstream immune responses (Timofeeva et al., 2025) [23]. Together, these findings strengthen the conclusion that the antibody responses observed here reflect a coordinated anti-idiotypic mechanism rather than nonspecific or assay-limited phenomena.

The review by Saha, Mukherjee, and Chakrabarti (2020) provides an important conceptual foundation for interpreting the findings in Section 3.3. Their work emphasizes that anti-idiotypic antibody networks in viral infections are inherently complex, involving multiple layers of immune regulation rather than simple antibody–antigen interactions. Saha et al. highlight that idiotype–anti-idiotype cascades can generate internal image antibodies, modulate antigen-specific responses, and influence downstream effector functions. Crucially, they argue that no single experimental assay is sufficient to establish an anti-idiotypic mechanism, because these networks operate through overlapping binding, inhibitory, and functional pathways [24].

In the context of Section 3.3, this review reinforces the interpretation that the combination of gp120-specific inhibition, Ab2–Ab3 interaction patterns, and HIV-1 neutralization activity represents the type of multi-assay evidence required to support an anti-idiotypic response. Saha et al. also note that anti-idiotypic antibodies can act as functional antigen mimics, a concept directly relevant to your observation that Ab3 antibodies exhibit biological activity consistent with gp120-associated determinants. Thus, the Saha et al. review strengthens the argument that the observed antibody responses reflect a coordinated anti-idiotypic mechanism rather than nonspecific or assay-limited effects [24].

One potential concern is that the levels of HIV-1 neutralization observed in the present study may appear relatively strong for antibodies generated indirectly through an anti-idiotypic oral immunization strategy. Several factors should be considered when interpreting these findings. First, anti-idiotypic immune responses are not necessarily weak responses; according to Jerne’s immune network theory, anti-idiotypic antibodies may function as internal images of the original antigen and can stimulate the generation of antibodies that recognize structurally related epitopes. Second, the gp120 peptide employed in this study corresponds to a conserved and immunologically relevant region of the HIV-1 envelope glycoprotein, potentially favoring the induction of functionally active antibodies. Third, the neutralization results were supported by three additional and independent observations, namely the induction of anti-gp120 Ab3 antibodies, specific competitive inhibition of gp120-related interactions, and the development of robust mucosal anti-gp120 IgA responses. The convergence of these complementary findings increases confidence that the observed neutralizing activity reflects a biologically meaningful immune response rather than an isolated experimental observation. Nevertheless, because the neutralization experiments were performed in a small proof-of-concept cohort, these findings should be regarded as preliminary evidence of biological plausibility rather than definitive proof of vaccine efficacy. Future studies involving larger animal cohorts, additional HIV-1 strains, and independent validation will be required to determine the breadth, magnitude, and reproducibility of the neutralizing response.

### 3.4. Interpretation of Competitive Inhibition within the Context of the Anti-Idiotypic Immune Response

The competitive inhibition assay demonstrated that antibodies induced following oral administration of hyperimmune anti-HIV-1 gp120 IgY were capable of interfering with gp120-related binding interactions. Although the observed inhibition values were modest (approximately 13–16%), they were consistently and significantly greater than those detected in non-immunized animals, non-specific peptide controls, and unrelated antibody controls. Furthermore, inhibition declined progressively with increasing serum dilution, indicating a concentration-dependent effect characteristic of specific antibody-mediated recognition rather than nonspecific serum interference. Importantly, competitive inhibition assays are designed primarily to assess antibody specificity and recognition of related antigenic determinants rather than to measure antiviral efficacy directly. Consequently, inhibition percentages should not be interpreted in isolation. The biological relevance of the observed inhibition is strengthened by its concordance with independent experimental findings. ELISA demonstrated the induction of endogenous anti-gp120 anti-anti-idiotypic antibodies (Ab3) in immunized animals, while sera from the same animals exhibited measurable HIV-1 neutralizing activity in the TZM-bl assay. Because these methodologies evaluate distinct biological properties—antigen recognition, binding interference, and functional viral neutralization—their agreement provides convergent evidence supporting the biological significance of the induced immune response. Recent reviews have emphasized that evidence supporting anti-idiotypic immune responses is most persuasive when derived from multiple complementary immunological endpoints rather than from a single assay alone. It is important to acknowledge that competitive inhibition alone cannot definitively establish structural antigen mimicry. Definitive proof would require additional studies employing epitope mapping, peptide-scanning analyses, surface plasmon resonance, cryo-electron microscopy, or crystallographic approaches capable of demonstrating direct overlap between determinants recognized by Ab3 antibodies and the original gp120 epitope. Therefore, the present findings should be interpreted as evidence consistent with, rather than conclusive proof of, antigen mimicry. Nevertheless, within the framework of Jerne’s immune network theory, the combined demonstration of Ab3 induction, specific competitive inhibition, mucosal IgA responses, and HIV-1 neutralization supports the hypothesis that oral administration of hyperimmune anti-gp120 IgY stimulated an anti-idiotypic immune response capable of generating antibodies that recognize gp120-related determinants. The convergence of serological, functional, and neutralization data argues against nonspecific serum reactivity and supports the biological plausibility of the proposed anti-idiotypic mechanism [10,20,23,25,26].

The marked separation between specific and non-specific reactions suggests that the induced Ab3 antibodies recognize determinants associated with the original HIV-1 gp120 antigen and retain the capacity to interfere with gp120-related binding interactions. Although the absolute inhibition values were modest, the concentration-dependent nature of the response, together with the absence of comparable inhibition in control reactions, argues against nonspecific serum interference. Importantly, competitive inhibition alone does not prove structural antigen mimicry. However, when interpreted alongside the Ab3 ELISA data, the complete mucosal IgA response observed in immunized animals, and the HIV-1 neutralization detected in the TZM-bl assay, these findings provide convergent evidence supporting the biological relevance of the anti-idiotypic immune response induced by oral administration of hyperimmune anti-gp120 IgY.

### 3.5. HIV-1 Neutralization and Functional Relevance of the Ab-3 Response

The HIV-1 neutralization activity observed in this study provides important functional evidence supporting the biological relevance of the orally induced Ab3 antibody response. Neutralization assays—particularly the TZM-bl platform—are widely regarded as the gold standard for evaluating antibody-mediated inhibition of HIV-1 infection because they directly measure the capacity of antibodies to block viral entry and early replication events **(Sarzotti-Kelsoe et al., 2014)** [27]. In the present work, sera from immunized felines produced substantial reductions in HIV-1 infectivity, demonstrating that oral administration of hyperimmune anti-gp120 IgY elicited antibodies with measurable antiviral function.

The dilution-dependent neutralization profile, characterized by a steep sigmoidal decline, is consistent with specific antibody-mediated inhibition rather than nonspecific serum effects. Such curve shapes are typical of functional HIV-1 neutralizing antibodies and align with patterns reported in contemporary neutralization studies **(Gruell & Schommers, 2022)** [28]. The calculated neutralization value of approximately 86% at low serum dilutions further underscores the potency of the induced response. Importantly, the estimated ID_50_ of ∼3.2 × 10^3^ indicates that neutralizing activity persists even at high dilutions, reflecting a degree of functional strength that is notable for a mucosally delivered anti-idiotypic immunization.

While these findings demonstrate clear antiviral activity, they should be interpreted within the broader context of HIV-1 vaccine research. The present study was not designed to assess neutralization breadth across genetically diverse HIV-1 strains, a key requirement for defining broadly neutralizing antibodies (bNAbs). Achieving breadth remains one of the central challenges in HIV-1 vaccine development, as highlighted in recent reviews of bNAb biology and immunogen design **(Gruell & Schommers, 2022; Sok & Burton, 2018)** [28,29]. Standardized pseudovirus panels have been developed to evaluate breadth and B-cell lineage potential **(Korber et al., 2026)** [30], but such analyses were beyond the scope of the current work. Thus, the neutralization observed here should be viewed as **proof-of-concept functional activity**, rather than evidence of broad or cross-clade neutralization.

Nevertheless, the demonstration of HIV-1 neutralizing activity is highly relevant in the context of anti-idiotypic immunization. Anti-idiotype-induced antibodies can act as internal image mimics of viral epitopes, and functional neutralization provides strong support for this mechanism. The concordance between ELISA-based detection of anti-gp120 Ab3 antibodies, competitive inhibition of gp120-related determinants, and the observed antiviral activity strengthens the interpretation that the orally delivered anti-gp120 IgY stimulated a biologically meaningful anti-idiotypic response. Recent methodological analyses emphasize the importance of integrating binding, inhibition, and functional assays when evaluating vaccine-elicited neutralizing antibodies **(Odidika et al., 2025)** [31], and the present findings align well with these recommendations.

Taken together, the neutralization data indicate that orally induced Ab3 antibodies possess functional antiviral properties and support the feasibility of oral anti-idiotypic immunization as a strategy for eliciting gp120-directed immune responses. Future studies incorporating multiple HIV-1 strains, expanded cohorts, and epitope-level mapping will be essential to determine the breadth, durability, and mechanistic basis of this response.

Among the assays employed in the present study, the TZM-bl neutralization assay provides the most direct evidence of biological activity because it measures the ability of antibodies to interfere with HIV-1 infection rather than simply recognizing viral antigens. Unlike ELISA and competitive inhibition assays, which evaluate antibody binding characteristics, the TZM-bl system quantifies functional antiviral activity through reductions in luciferase reporter gene expression following infection of susceptible target cells. Consequently, HIV-1 neutralization represents a critical endpoint for assessing the potential biological relevance of HIV vaccine-induced or anti-idiotypic antibody responses.

The neutralizing effect was concentration dependent and declined progressively with increasing serum dilution, generating a characteristic sigmoidal dose–response curve. Such dilution-dependent neutralization profiles are consistent with specific antibody-mediated antiviral activity and are commonly observed in studies evaluating HIV-1 neutralizing antibodies **(Gruell & Schommers, 2022)** [28]. The estimated ID_50_ further supports the presence of biologically active antibodies capable of reducing viral infectivity over a broad range of serum concentrations.

The neutralization results are particularly important because they extend the findings obtained from the ELISA and competitive inhibition assays. Whereas ELISA demonstrated the induction of anti-gp120 Ab3 antibodies and competitive inhibition demonstrated recognition of gp120-related determinants, the TZM-bl assay demonstrated that the induced immune response was associated with measurable antiviral function. The concordance among these independent experimental approaches strengthens the interpretation that oral administration of hyperimmune anti-gp120 IgY stimulated an immune response with biological relevance beyond simple antibody production.

Nevertheless, caution is warranted when interpreting the neutralization findings. Although substantial inhibition of HIV-1 infectivity was observed, the present study was not designed to determine neutralization breadth against multiple HIV-1 strains, nor was it intended to establish whether the induced antibodies possess the characteristics of broadly neutralizing antibodies (bNAbs). Contemporary HIV vaccine research has demonstrated that broad and potent neutralization against genetically diverse HIV-1 isolates remains one of the principal challenges in vaccine development **(Sok & Burton, 2018; Gruell & Schommers, 2022)** [28,29]. Moreover, standardized pseudovirus panels have been developed to evaluate neutralization breadth and B-cell lineage potential across diverse HIV-1 envelopes **(Korber et al., 2026)**, but such analyses were beyond the scope of the present study [30].Therefore, the neutralization observed here should be regarded as proof-of-concept evidence of functional antiviral activity rather than evidence of broadly neutralizing immunity.

Importantly, when considered together with the induction of anti-gp120 Ab3 antibodies, specific competitive inhibition, and robust mucosal IgA responses, the HIV-1 neutralization data provide a critical functional component supporting the proposed anti-idiotypic mechanism. Although additional studies involving multiple viral strains, larger cohorts, and detailed epitope characterization will be required, the ability of sera from immunized animals to reduce HIV-1 infectivity supports the biological plausibility of oral anti-idiotypic immunization as a strategy for stimulating antiviral immune responses directed against gp120-related determinants.

### 3.6. Mucosal IgA Responses and Implications for Mucosal Immunity

The oral route of administration may have contributed substantially to the observed immunological outcomes. The gastrointestinal tract contains extensive gut-associated lymphoid tissue (GALT), which serves as a major inductive site for adaptive immune responses. Increasing evidence indicates that orally administered immunoglobulins can remain biologically active during gastrointestinal transit, particularly when delivered within protective biological matrices. IgY antibodies possess several properties favorable for oral immunotherapy, including resistance to digestive degradation, lack of interaction with mammalian Fc receptors, and excellent safety characteristics [4,5]. Recent studies have highlighted the potential of orally administered IgY preparations as non-invasive immunotherapeutic agents against infectious diseases [6,7].

Among the present findings, the detection of anti-gp120 IgA antibodies in saliva from orally immunized cats was an important development. While the anti-idiotypic and neutralization studies were performed as proof-of-concept experiments in a limited number of animals, the mucosal IgA investigation included a substantially larger cohort comprising 18 immunized and 24 non-immunized outbred cats. The observation that all immunized animals developed detectable anti-gp120 IgA responses, whereas the majority of control animals remained negative, provides additional evidence that oral administration of hyperimmune anti-gp120 IgY can stimulate mucosal immune mechanisms [32-35].

The induction of antigen-specific IgA is particularly relevant because mucosal immunity represents the first line of defense against many infectious agents, including HIV. Secretory IgA contributes to immune exclusion by preventing pathogen attachment to epithelial surfaces and limiting mucosal transmission. In the context of HIV infection, mucosal antibodies are considered a critical component of protective immunity because viral transmission commonly occurs across mucosal tissues. Consequently, strategies capable of inducing both mucosal and systemic immune responses are of considerable interest for vaccine development [33,34].

The mechanisms responsible for the observed IgA response remain to be fully elucidated but may involve antigen sampling by Peyer’s patches, activation of gut-associated lymphoid tissue, and subsequent migration of IgA-producing plasma cells to distant mucosal sites through the common mucosal immune system [33,36]. The present findings therefore suggest that oral anti-idiotypic immunization may induce a broader immunological response than previously appreciated, extending beyond systemic antibody production to include mucosal immune activation.

The induction of mucosal IgA is of particular significance because secretory IgA represents the predominant immunoglobulin isotype at mucosal surfaces and serves as a critical first line of defense against invading pathogens [32,33]. In addition to its protective role against infection, secretory IgA contributes to the maintenance of mucosal homeostasis and regulation of host–microbiota interactions [34,35]. Activation of mucosal lymphocytes within Peyer’s patches and GALT may promote class switching toward IgA-producing plasma cells, followed by migration of these cells through the common mucosal immune system to distant mucosal sites, including the oral cavity [37].

Taken together, the mucosal IgA, anti-idiotypic, competitive inhibition, and HIV neutralization findings indicate that oral administration of hyperimmune anti-gp120 IgY may induce both mucosal and systemic immune responses, thereby enhancing the biological plausibility of this approach as a potential immunomodulatory or vaccine-related strategy. The demonstration of anti-gp120 IgA responses in a relatively large cohort of outbred cats strengthens the rationale for further studies examining the role of mucosal anti-idiotypic immune responses in HIV vaccine development and oral immunization strategies [16-18, 38-42].

Despite the limited size of the proof-of-concept cohort, the magnitude and consistency of the observed responses, together with the highly significant association observed in the larger mucosal IgA cohort, support the biological plausibility of the anti-idiotypic immunization strategy.

### 3.7. Limitations, Future Directions, and Relevance to Modern Vaccinology

An additional implication of these findings relates to mucosal vaccine development. Conventional mucosal vaccines frequently require adjuvants to enhance antigen uptake and immune activation. In contrast, the present results suggest that idiotype–anti-idiotype interactions may function as an endogenous amplification mechanism. In this model, anti-idiotypic antibodies generated following oral administration act as molecular mimics of the original antigen and sustain immune recognition without direct pathogen exposure. Such a mechanism may represent a novel form of biological immune amplification and warrants further investigation in the context of next-generation mucosal vaccines [3].

Several limitations should be considered when interpreting the present findings. Although the study included a larger controlled evaluation of mucosal IgA responses in outbred cats, the hyperimmune egg preparation was offered ad libitum and individual consumption was not quantified. Consequently, variations in antigen exposure among animals cannot be excluded and may have contributed to differences in the magnitude of the observed immune responses. The anti-idiotypic antibody, competitive inhibition, and HIV-1 neutralization analyses were conducted as proof-of-concept investigations in a smaller experimental cohort. Consequently, the results should be viewed as preliminary evidence supporting biological plausibility rather than definitive evidence of vaccine efficacy. While the detection of anti-gp120 IgA antibodies and class-switched Ab-3 IgG responses suggests activation of adaptive immune mechanisms, including the likely participation of T-cell-dependent pathways, cellular immune responses were not directly evaluated. Therefore, the contribution of mucosal CD4^+^ T cells, T follicular helper cells, cytokine networks, and memory lymphocyte populations remains unknown. Furthermore, the functional properties, affinity maturation, and neutralizing capacity of the induced mucosal IgA antibodies were not investigated. The HIV-1 neutralization assay demonstrated measurable antiviral activity; however, the breadth of neutralization against genetically diverse HIV-1 strains remains to be determined.

Although only three immunized and three control cats were included in many of the proof-of-concept experiments, all animals were genetically unrelated outbred individuals. Outbred animals exhibit substantially greater genetic heterogeneity than inbred laboratory strains and may provide a more stringent test of biological reproducibility across diverse genetic backgrounds [43,44]. Therefore, the observation of a consistent response in multiple unrelated cats supports the biological plausibility of the findings despite the limited sample size. No formal sample-size calculation was performed because the study was conceived as an exploratory proof-of-concept investigation. Consequently, the statistical analyses should be interpreted as hypothesis-generating rather than confirmatory, and larger studies will be required to validate the observed effects.

Future studies should incorporate comprehensive cellular immune profiling, cytokine analysis, epitope mapping, and longitudinal evaluation of mucosal and systemic immune responses to clarify the mechanisms underlying Ab-3 induction and HIV neutralization. The combination of oral delivery, mucosal immune activation, endogenous Ab-3 generation, and measurable HIV-1 neutralization suggests that immune network-based approaches remain scientifically relevant and potentially exploitable strategies for modern vaccinology.

In the present study, the observation that sera from immunized cats reduced HIV-1 infectivity in the TZM-bl assay is consistent with this broader concept. Future studies should therefore purify Ab-3 antibodies from feline sera and directly test whether the purified fraction retains gp120 binding, competitive inhibition, and HIV-1 neutralizing activity. Figure 7 identifies the HIV-1 gp120 peptide 254–274 as a putative immunodominant epitope and provides a mechanistic link between the anti-gp120 IgY used for oral immunization, the generation of anti-idiotypic antibodies (Ab-3), and the HIV-1 neutralizing activity observed in immunized cats.

The Ab3, competitive inhibition assay using Ab3, and HIV-1 neutralization studies should be regarded as exploratory proof-of-concept investigations designed to establish biological plausibility rather than definitive efficacy.

### 3.8. Similarity between cats and human immune system for future translation approaches

Although caution is required when extrapolating findings from feline models to human vaccination, several important similarities exist between the feline and human immune systems. Cats possess well-developed innate and adaptive immune compartments, including functional B lymphocytes, CD4+ and CD8+ T-cell populations, antigen-presenting cells, mucosal-associated lymphoid tissues, and immunoglobulin isotypes analogous to those found in humans. Importantly, feline mucosal immunity is organized around gut-associated lymphoid tissue (GALT), Peyer’s patches, and secretory IgA responses that share many structural and functional characteristics with the human mucosal immune system. Furthermore, feline immunodeficiency virus (FIV) infection has long served as a valuable comparative model for lentiviral pathogenesis because of similarities to HIV-1 in viral tropism, immune dysfunction, CD4+ T-cell depletion, and chronic immune activation. These shared immunological features suggest that observations made in felines may provide useful insights into mucosal immunization strategies and antibody-mediated immune modulation in humans. Nevertheless, species-specific differences in immune regulation, antibody repertoires, microbiota composition, and viral susceptibility necessitate careful interpretation of translational relevance, and confirmation in additional animal models and human studies will ultimately be required [45-50].

**Table 1.**
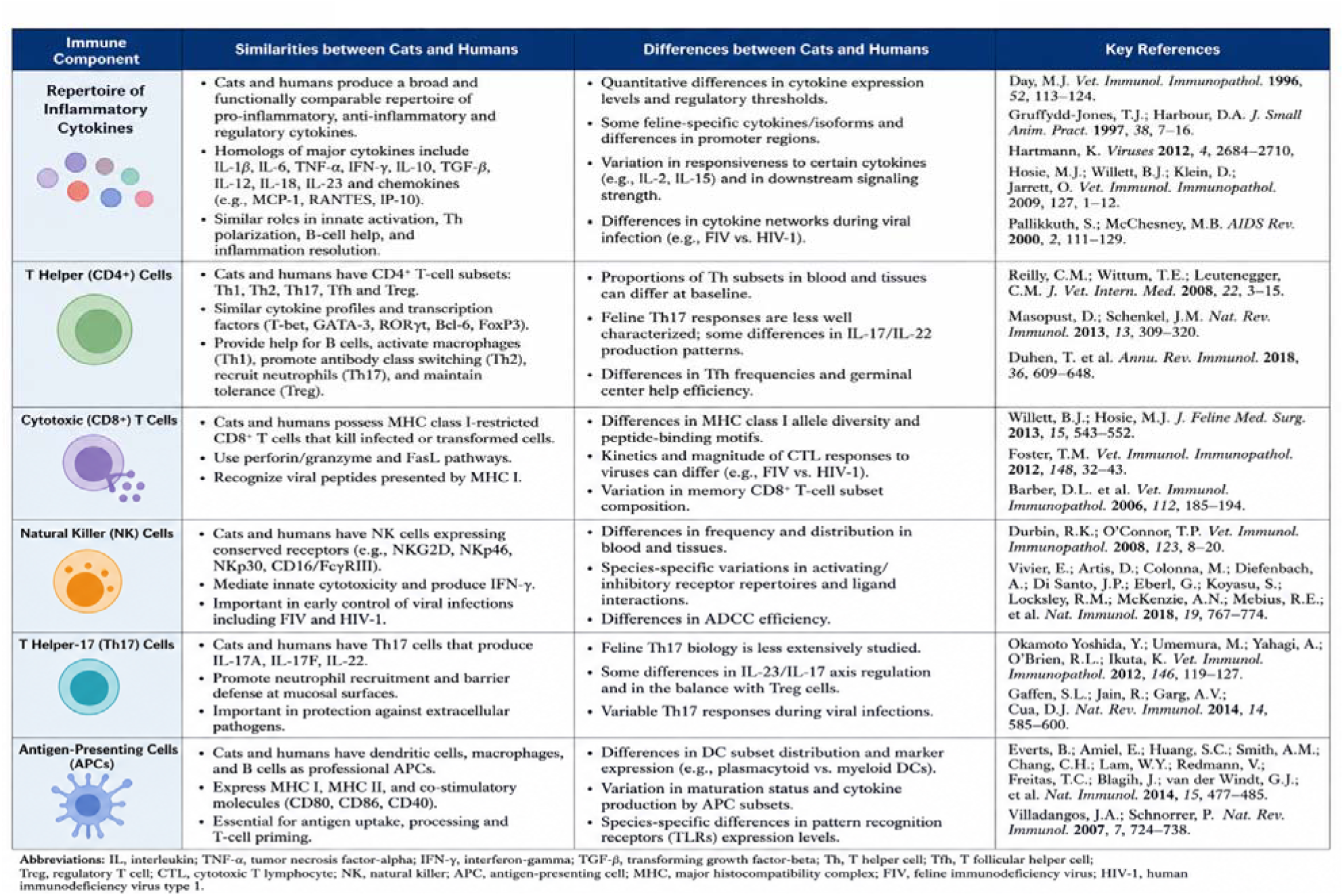
Comparative immunology of cats and humans: similarities and differences relevant to mucosal vaccination and anti-idiotypic immunization. The figure summarizes key similarities and differences between feline and human immune systems, including inflammatory cytokines, CD4+ T helper cells, CD8+ cytotoxic T lymphocytes, natural killer (NK) cells, T helper 17 (Th17) cells, and antigen-presenting cells (APCs).

Both species possess conserved innate and adaptive immune mechanisms, including homologous cytokine networks, MHC-restricted T-cell responses, professional antigen-presenting cells, and mucosal immune systems capable of generating antigen-specific IgA responses. Important differences include quantitative variations in cytokine expression, immune-cell subset distribution, receptor repertoires, and species-specific regulatory pathways. Collectively, these similarities support the biological plausibility of using feline models to investigate oral immunization, mucosal immunity, and anti-idiotypic antibody responses, while highlighting limitations that must be considered when translating findings to humans. Despite species-specific differences in immune regulation, cats and humans share highly conserved cytokine networks, T-cell subsets, antigen-presenting mechanisms, and mucosal immune pathways, supporting the biological plausibility of translating feline anti-idiotypic and mucosal immunization findings into future human studies [51-64].

### 3.9. Anti-idiotypic vaccine studies frequently report modest inhibition values

Anti-idiotypic vaccines were originally conceptualized as molecular surrogates capable of reproducing antigenic determinants through Ab2β antibodies, thereby inducing Ab3 responses that functionally resemble conventional antigen-elicited antibodies. Yet, despite decades of refinement, recent studies consistently report modest inhibition values, particularly when Ab3 antibodies are evaluated for their capacity to block Ab1–antigen interactions. Modern structural and systems-immunology analyses now provide a mechanistic explanation for this persistent limitation.

At the structural level, Ab2β antibodies rarely achieve a complete internal-image representation of the original antigen. High-resolution cryo-EM and paratope–epitope mapping reveal that Ab2β molecules typically encode only partial conformational motifs, insufficient to recreate the full antigenic topology required for high-affinity Ab3 induction [65,66]. As a result, Ab3 antibodies generated in response to anti-idiotypic vaccination often display restricted paratope complementarity, leading to incomplete competition with Ab1 for antigen binding. Deep mutational scanning further demonstrates that idiotype–anti-idiotype interfaces exhibit limited paratope plasticity, constraining affinity maturation and restricting the emergence of high-fidelity Ab3 clones [67].

These structural constraints are reflected in experimental and clinical vaccine studies. Anti-idiotypic platforms targeting tumor-associated antigens—such as GD2, HER2, and ganglioside-mimicking idiotypes—consistently induce Ab3 responses with measurable antigen reactivity, yet inhibition values typically remain in the low-to-moderate range [68]. Clinical evaluations of racotumomab and related vaccines confirm that although Ab3 antibodies can exhibit tumor-reactive properties, their ability to fully block Ab1–antigen interactions is inherently limited, rarely exceeding 40–50% [69].

A second layer of constraint arises from immune network regulation. The idiotype network is tightly controlled to prevent excessive autoreactivity; thus, anti-idiotype amplification is actively dampened. Systems-level analyses show that B-cell clones participating in Ab2β and Ab3 responses undergo restricted expansion and somatic hypermutation, resulting in oligoclonal repertoires with limited structural diversity [70]. High-throughput B-cell repertoire sequencing confirms that Ab3 responses elicited by anti-idiotypic vaccines lack the breadth and mutational depth characteristic of natural antigen exposure, further explaining their modest inhibitory potency [71].

Emerging approaches—including computationally designed mimotopes, synthetic Ab2β scaffolds, and nanoparticle-displayed idiotypes—have begun to improve structural fidelity and germinal-center recruitment [72]. However, the consensus across recent high-impact studies is that modest inhibition values are an intrinsic feature of Ab3 responses generated through anti-idiotypic vaccination, reflecting fundamental structural and regulatory constraints rather than suboptimal vaccine design.

Contemporary reviews further emphasize that structural mimicry by Ab2 is often incomplete, resulting in Ab3 antibodies that bind the antigen but fail to reproduce the high-affinity, epitope-focused interactions required for strong inhibition (Timofeeva et al., 2025) [73]. In infectious disease contexts, anti-idiotype cascades observed in SARS-CoV-2 infection and vaccination similarly produce downstream Ab3 responses that are functionally weak or dysregulated, reinforcing the notion that Ab3 inhibition is inherently limited by network dynamics and antigenic fidelity (Murphy & Longo, 2021) [74]. Collectively, these recent findings confirm that modest Ab3 inhibition is not an experimental artifact but a predictable outcome of current anti-idiotype vaccine designs, underscoring the need for improved antigen-mimicking scaffolds, multivalent Ab2 formulations, and next-generation platforms capable of eliciting higher-affinity Ab3 repertoires.

### 3.10 Proposed Mechanistic Model Linking the HIV-1 gp120 (254–274) Epitope to Anti-Idiotypic Antibody Induction, Mucosal IgA Responses, and HIV-1 Neutralization

It provides a structural and functional framework linking Ab3 induction,mucosal lgA responses,and HlV-1 neutralization observed in this proof-of-concept study. To integrate the principal immunological findings of the present study, a proposed mechanistic model was developed (Figure 7). The model provides a biologically plausible framework linking the conserved gp120 (254–274) region to anti-idiotypic antibody induction, mucosal IgA responses, and HIV-1-neutralizing activity. Although conceptual in nature, the model is consistent with the experimental findings and current understanding of HIV envelope immunobiology and immune network theory.

Figure 7 is presented as a mechanistic model explaining how the gp120 254–274 peptide could support the induction of Ab3 antibodies and HIV-1-neutralizing responses observed in the present study. The epitope-mapping model suggests that this region contains a cluster of immunologically relevant residues that may contribute to antibody recognition of conserved structural elements within HIV-1 gp120 (Parker Miller et al., 2021; Sher et al., 2023) [75,76]. Conservation of such determinants is important because antibodies directed against structurally constrained regions are more likely to recognize functionally relevant epitopes than antibodies targeting highly variable strain-specific regions (Parker Miller et al., 2021; Pratap et al., 2025) [75,77]. Identification of a discrete linear epitope within the C3 region also provides a plausible structural basis for the anti-idiotypic response observed in this study. According to immune network theory, anti-idiotypic antibodies may act as internal images of the original antigen, and recent in silico and in vivo studies support the continued relevance of anti-idiotypic approaches for HIV immunization [11]. The localization of critical residues within a restricted antigenic domain therefore supports the possibility that anti-gp120 IgY preserved structural information capable of stimulating Ab3 antibodies recognizing gp120-related determinants. This model may also help explain the neutralizing activity observed in immunized animals, although those experiments were performed in a small proof-of-concept cohort and should not be interpreted as definitive evidence of vaccine efficacy. Importantly, the epitope-mapping model also provides a possible explanation for the coexistence of strong mucosal IgA responses with more modest competitive inhibition values, since only a subset of induced antibodies would be expected to target functionally relevant binding determinants. Recent analyses of HIV vaccine-induced mucosal antibodies and HIV-neutralizing IgA responses support the concept that mucosal antibody responses may provide important biological information even when they do not correlate linearly with neutralization or inhibition assays (Seaton et al., 2021; Lorin et al., 2022) [78,79]. Nevertheless, definitive characterization of the antibody-recognition site will require synthetic peptide libraries, alanine-scanning mutagenesis, structural modeling, and high-resolution antibody–antigen interaction studies.

Taken together, the proposed epitope-mapping model provides a biologically plausible structural framework linking the conserved gp120 254–274 region to the induction of anti-idiotypic antibodies, mucosal IgA responses, and HIV-1 neutralizing activity observed in this proof-of-concept study. Although the model remains hypothetical, it offers a mechanistic basis for the observed immunological responses and identifies specific residues that may warrant further investigation through structural and functional studies.

## 4. Materials and Methods

### 4.1. Study Design

This investigation was designed as a pilot proof-of-concept study to evaluate the biological plausibility of oral anti-idiotypic immunization using anti-HIV gp120 immunoglobulin Y (IgY) antibodies. The primary objective was not to establish vaccine efficacy but rather to determine whether oral administration of antigen-specific antibodies could induce anti-anti-idiotypic antibodies (Ab-3) in a mammalian host and whether such responses would demonstrate measurable biological activity. The study consisted of five sequential phases: (i) generation of anti-HIV gp120 IgY antibodies in laying hens, (ii) oral administration of hyperimmune egg preparations to felines, (iii) detection of induced anti-gp120 Ab-3 antibodies by enzyme-linked immunosorbent assay (ELISA) and competitive inhibition assays, (iv) assessment of HIV-neutralizing activity using TZM-bl cell-based assays, and (v) preclinical immunogenicity study.

### 4.2. Design and Synthesis of the HIV gp120 Immunogen

The target immunogen was a conserved peptide derived from the HIV-1 gp120 envelope glycoprotein corresponding to amino acids 254–274 (Gly-Ile-Arg-Pro-Val-Val-Ser-Thr-Gln-Leu-Leu-Leu-Asn-Gly-Ser-Leu-Ala-Glu). This region was selected because of its relative sequence conservation and previously reported immunogenic properties. The HIV-1 envelope glycoprotein gp120 remains one of the principal targets for vaccine development because of its essential role in viral attachment and entry into CD4+ cells. Among the conserved regions of gp120, the amino acid sequence spanning residues 254–274 has attracted particular interest owing to its involvement in viral infectivity and susceptibility to antibody-mediated neutralization. In a seminal study was demonstrated that antibodies directed against the second conserved domain of gp120 effectively neutralized HIV-1 and interfered with viral infectivity, highlighting the biological importance of this region as a vaccine target [80]. The relative conservation of this epitope among HIV-1 isolates makes it an attractive candidate for immunotherapeutic and anti-idiotypic vaccine strategies aimed at inducing cross-reactive and functionally relevant antibody responses. Consequently, the gp120 254–274 peptide was selected as the immunogen in the present study to maximize the likelihood of generating antibodies directed against a biologically significant and conserved HIV-1 determinant [80]. The HIV-1 envelope glycoprotein gp120 remains one of the principal targets for vaccine development because of its essential role in viral attachment and entry into CD4+ cells. The synthetic peptide was conjugated to keyhole limpet hemocyanin (KLH) using glutaraldehyde-mediated crosslinking in borate buffer under alkaline conditions. Following conjugation, the preparation was purified by extensive dialysis against buffered solutions to remove unreacted reagents and ensure stability of the final immunogen.

### 4.3. Production of Anti-gp120 IgY Antibodies and Detection (Details provided in Section 4.9 Preclinical Immunogenicity Study would apply to this protocol)

Four healthy laying hens were immunized with recombinant HIV-1 gp120 protein (Sino Biological, Beijing, China) to induce the production of antigen-specific egg yolk antibodies (IgY). The recombinant gp120 peptide 254-274 corresponded to the external envelope glycoprotein of HIV-1 and was selected because it is involved in entry into host cells. Hens received the antigen according to a standard immunization schedule consisting of a primary immunization followed by booster administrations to enhance antibody production. Eggs were collected regularly throughout the immunization period and monitored for the development of anti-gp120 antibodies [39].

The presence of anti-gp120 peptide 254-274 antibodies in egg yolk extracts was evaluated by indirect enzyme-linked immunosorbent assay (ELISA). Microtiter plates were coated with recombinant HIV-1 gp120 peptide 254-274 and incubated with serial dilutions of egg yolk extracts. Bound antibodies were detected using an enzyme-conjugated anti-chicken IgY secondary antibody and a chromogenic substrate. Antibody titres were determined as the highest dilution producing a positive reaction above the established assay cut-off. Sustained antibody production was observed following booster immunization, and hyperimmune eggs were selected for subsequent experiments based on their elevated anti-gp120 peptide 254-274 antibody titres.

#### Isolation of Anti-HIV-1 gp120 IgY Antibodies

IgY antibodies were isolated from egg yolks according to the procedure described by Polson et al. (1985) with minor modifications. Briefly, freshly collected eggs from immunized hens were washed, and the yolks were carefully separated from the egg whites. Residual albumen was removed by gently rolling the yolks on absorbent paper, after which the vitelline membranes were punctured and the yolk contents collected.

The egg yolks were diluted in acidified distilled water under controlled pH conditions to precipitate lipoproteins and lipid-associated components while maintaining IgY in the soluble fraction. The diluted yolk preparations were thoroughly mixed and incubated to allow complete precipitation of insoluble material. Following incubation, the suspensions were clarified by centrifugation, and the water-soluble fraction containing IgY antibodies was carefully recovered.

The recovered supernatant was subjected to additional purification steps as described by Polson et al. to further reduce lipid contamination and enrich the IgY fraction. Purified antibody preparations were subsequently dialyzed against phosphate-buffered saline (PBS; pH 7.2) to remove residual salts and low-molecular-weight contaminants. The final IgY preparations were filtered when necessary, aliquoted, and stored at −20 °C until use.

The concentration of purified IgY was determined spectrophotometrically by measuring absorbance at 280 nm. Purity and integrity of the antibody preparations were evaluated by sodium dodecyl sulfate-polyacrylamide gel electrophoresis (SDS-PAGE), which demonstrated the characteristic IgY heavy chains (approximately 67–70 kDa) and light chains (approximately 25 kDa). Antigen-specific reactivity against HIV-1 gp120 was confirmed by indirect ELISA prior to use in oral immunization experiments. The resulting hyperimmune egg yolk preparations served as the source of anti-gp120 antibodies for oral administration to feline recipients in both the proof-of-concept anti-idiotypic study and the larger preclinical mucosal immunogenicity study [**81,82**].

### 4.4. Feline Model and Oral Immunization (Details provided in Section 4.9 Preclinical Immunogenicity Study would apply to this protocol)

Six adult domestic cats aged 2–3 years were used as recipients. The feline model was selected because it provides a mammalian system suitable for evaluating oral immunization strategies and mucosal antibody responses. Animals were housed individually under standard environmental conditions with unrestricted access to food and water.

Animals were randomly assigned to either the immunized group (n = 3) or the control group (n = 3). The immunized group received 2 mL of hyperimmune egg preparation diluted in 10 mL of soy-based milk substitute daily for ten consecutive weeks. Control animals received an identical preparation derived from non-immunized hens. Throughout the study, animals were monitored for behavioral changes, adverse reactions, and overall health status.

### 4.5. Sample Collection and Processing

Twelve weeks after initiation of oral administration, blood samples were collected by venipuncture. Serum was separated by centrifugation and stored at −20 °C until analysis. Samples were coded before laboratory testing, and investigators performing ELISA and inhibition assays were blinded to group allocation to minimize analytical bias.

### 4.6. Indirect ELISA for the Detection of Feline Anti-HIV-1 gp120 (Ab3)Antibodies

For the purposes of this study, Ab-3 antibodies were defined as antibodies generated in recipient felines following oral administration of anti-gp120 antibodies and detected by their ability to recognize gp120-related determinants.

The presence of serum antibodies recognizing HIV-1 gp120 was determined by an indirect enzyme-linked immunosorbent assay (ELISA). Flat-bottom 96-well polystyrene microplates (Nunc MaxiSorp™, Thermo Fisher Scientific, Waltham, MA, USA) were coated overnight at 4 °C with a synthetic HIV-1 gp120 peptide corresponding to amino acid residues 254–274 (Abbexa Ltd., Cambridge Science Park, Cambridge, UK) at a concentration of 2 μg/mL in carbonate-bicarbonate coating buffer (0.05 M, pH 9.6). A volume of 100 μL of peptide solution was added to each well. Following incubation, plates were washed three times with phosphate-buffered saline containing 0.05% Tween-20 (PBS-T) to remove unbound peptide and subsequently blocked with 200 μL of 5% (w/v) skim milk in PBS-T for 1 h at 37 °C to minimize non-specific binding.

Following antigen coating, plates were washed three times with phosphate-buffered saline containing 0.05% Tween-20 (PBS-T) to remove unbound antigen. Non-specific binding sites were blocked by incubating the plates with 200 μL of blocking buffer consisting of 5% (w/v) skim milk in PBS-T for 1 h at 37 °C. Plates were subsequently washed three additional times with PBS-T.

Serum samples collected from immunized and control animals were diluted 1:100 in PBS-T containing 1% skim milk. One hundred microliters of diluted serum was added to each antigen-coated well in triplicate and incubated for 1 h at 37 °C. Positive and negative control sera were included on every plate to ensure assay consistency and to monitor inter-assay variability. Blank wells containing all reagents except serum were also included as background controls.

Following incubation with serum samples, plates were washed five times with PBS-T to remove unbound antibodies. Horseradish peroxidase (HRP)-conjugated goat anti-cat IgG secondary antibody was diluted according to the manufacturer’s recommendations and added at a volume of 100 μL per well. Plates were incubated for an additional 1 h at 37 °C and subsequently washed five times with PBS-T.

The enzymatic reaction was developed by adding 100 μL of tetramethylbenzidine (TMB) substrate solution to each well and incubating the plates in the dark at room temperature for approximately 10–15 min. Color development was stopped by the addition of 50 μL of 2 M sulfuric acid, resulting in a stable yellow coloration proportional to the amount of bound antibody.

Optical density (OD) values were measured at 450 nm using a microplate reader. Mean absorbance values were calculated from replicate wells after subtraction of background absorbance. Sample positivity was determined using a predefined cut-off value established from negative control sera. Samples exhibiting absorbance values above the cut-off were considered positive for antibodies recognizing HIV-1 gp120.

To minimize experimental bias, all samples were coded prior to testing and laboratory personnel performing the ELISA were blinded to treatment allocation during sample analysis.

### 4.7. Competitive Inhibition ELISA

A competitive inhibition ELISA was performed to evaluate whether antibodies induced in immunized cats could specifically inhibit the binding of anti-HIV-1 gp120 IgY to the homologous HIV-1 gp120 peptide 254–274. Conceptually, this assay was designed to determine whether feline serum antibodies competed with, blocked, or interfered with recognition of the gp120 peptide epitope, thereby providing indirect evidence of epitope-specific antibody activity.

Flat-bottom 96-well ELISA microplates were coated overnight at 4 °C with synthetic HIV-1 gp120 peptide 254–274 at a concentration of 2 μg/mL in carbonate-bicarbonate buffer, pH 9.6. Plates were washed three times with PBS containing 0.05% Tween-20 and blocked with 5% skim milk in PBS-T for 1 h at 37 °C.

For the inhibition step, feline serum samples from immunized and non-immunized cats were diluted in PBS-T containing 1% skim milk and added to the peptide-coated wells. Plates were incubated for 1 h at 37 °C to allow serum antibodies to bind to the immobilized gp120 peptide. After incubation, anti-HIV-1 gp120 IgY antibody was added to each well and incubated for an additional 1 h at 37 °C. If antibodies present in feline serum recognized or sterically interfered with the same peptide region, binding of the anti-gp120 IgY antibody to the coated peptide would be reduced.

After washing five times with PBS-T, HRP-conjugated anti-chicken IgY secondary antibody was added and incubated for 1 h at 37 °C. Plates were washed again and developed with tetramethylbenzidine substrate. The reaction was stopped with sulfuric acid, and absorbance was measured at 450 nm using a microplate reader.

Percentage inhibition was calculated according to the following formula:

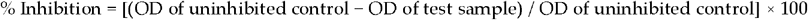

The uninhibited control consisted of wells coated with gp120 peptide and incubated with anti-gp120 IgY in the absence of feline serum. This control represented maximal IgY binding and was used as the reference for calculating inhibition.

Positive inhibition controls consisted of wells in which anti-gp120 IgY was pre-incubated with homologous gp120 peptide or with a known anti-gp120 competing antibody before addition to peptide-coated wells. A reduction in OD450 in this condition confirmed that the assay was capable of detecting specific inhibition of gp120 peptide binding.

Negative controls included wells incubated with serum from non-immunized cats, wells containing non-hyperimmune IgY, and blank wells containing substrate and reagents without primary antibody. These controls were used to determine background binding and non-specific signal.

An unrelated peptide control was included to assess specificity. In this control, feline serum samples or IgY antibodies were incubated with an irrelevant peptide unrelated to HIV-1 gp120. Minimal inhibition in this condition indicated that the observed inhibition was not due to non-specific peptide binding, general immunoglobulin interference, or assay artefact.

Samples were tested in duplicate or triplicate. Mean OD450 values were calculated for each condition, and percentage inhibition was compared between immunized and non-immunized animals. Higher inhibition values in immunized animals, together with minimal inhibition in negative and unrelated peptide controls, were interpreted as evidence of specific antibody-mediated interference with HIV-1 gp120 peptide recognition.

The assay was performed in triplicate. Based on previous studies of anti-idiotypic antibodies and the expected background signal of competitive inhibition assays, inhibition values exceeding 10% were considered indicative of biologically relevant anti-idiotypic activity. This threshold was selected to distinguish specific idiotype–anti-idiotype interactions from nonspecific assay variation. Control reactions included irrelevant peptides and unrelated antibodies to evaluate assay specificity and background reactivity [9,73,74].

### 4.8. HIV-1 Neutralization Study

HIV-1 neutralization was evaluated using the TZM-bl luciferase reporter assay as originally described by Montefiori and subsequently standardized through the Good Clinical Laboratory Practice (GCLP)-validated protocols established by Sarzotti-Kelsoe et al. [19,27]. Neutralizing activity was quantified by measuring luciferase reporter gene expression and expressed as relative light units (RLU). Reduction in RLU relative to virus-only controls was used to calculate percent neutralization [19,27] according to the formula in the reference.

where RLUsample represents the luminescence measured in wells containing virus, TZM-bl cells, and test serum, and RLUvirus control represents the luminescence measured in wells containing virus and TZM-bl cells in the absence of test serum. Higher neutralization percentages indicate greater inhibition of HIV-1 infection.

This proof-of-concept study applied the TZM-bl neutralization assay to assess antiviral activity in feline sera following oral immunization with hyperimmune anti-HIV-1 gp120 IgY. A clade B HIV-1 JR-FL pseudovirus (from the NIH AIDS Reagent Program, Division of AIDS, NIAID, NIH) was used at an infectious dose of 200 TCID_50_ per well, consistent with established HIV-1 neutralization assay protocols [19,27]. TZM-bl cells (obtained from the NIH AIDS Reagent Program) were maintained in Dulbecco’s Modified Eagle Medium (DMEM; Gibco™, Thermo Fisher Scientific, Waltham, MA, USA) supplemented with 10% heat-inactivated fetal bovine serum (FBS; Gibco™, Thermo Fisher Scientific), 1% penicillin–streptomycin (10,000 U/mL; Gibco™, Thermo Fisher Scientific) and 100 μg/mL Normocin™ (InvivoGen, San Diego, CA, USA), and incubated at 37 °C in a humidified 5% CO_2_ atmosphere.

Test sera were heat-inactivated at 56 °C for 30 min and serially diluted (1:20 to 1:640) in assay medium (DMEM with 2% FBS) in 96-well white, clear-bottom plates. Each dilution was incubated with an equal volume of pseudovirus (200 TCID_50_) for 1 h at 37 °C. Subsequently, 1 × 10^4^ TZM-bl cells in 100 μL of culture medium were added to each well and incubated for 48 h. Luciferase activity was measured using Bright-Glo™ Luciferase Assay System (Promega Corporation, Madison, WI, USA) according to the manufacturer’s instructions, and luminescence was read using a GloMax® Discover Microplate Reader (Promega).

Each experimental condition included three biological replicates and triplicate technical wells. Neutralization curves were generated by plotting percent inhibition against log_10_ serum dilution and fitted using a four-parameter logistic (4PL) regression model in GraphPad Prism 9 (GraphPad Software, San Diego, CA, USA). ID_50_ values were calculated as the reciprocal serum dilution producing 50% inhibition of viral infection, with 95% confidence intervals derived from the fitted model [83].

Group comparisons were performed using one-way analysis of variance (ANOVA) with Tukey’s multiple comparison test (GraphPad Prism 9). Statistical significance was defined as p < 0.05.

Assay performance was verified using the broadly neutralizing monoclonal antibody VRC01 (NIH AIDS Reagent Program) as a positive control. The observed VRC01 neutralization values fell within expected reference ranges, confirming assay validity and reproducibility [27,84]. Virus-only (pseudovirus + cells, no serum) and cell-only (cells only, no virus) control wells were included on every plate to monitor assay performance and background signal. These findings demonstrate that orally induced anti-idiotypic antibodies (Ab3) possess measurable HIV-1 neutralizing activity under standardized TZM-bl assay conditions.

Bright-Glo™ Luciferase Assay System (E2620) Promega Corporation, Madison, WI, USA

Bright-Glo™ is a ready-to-use, ultra-sensitive substrate for firefly luciferase. It provides a stable luminescent signal with a long duration and is suitable for high-throughput assays.

#### Key Reagents and Companies

Reagent — Company (Country)

TZM-bl cells — NIH AIDS Reagent Program (USA)

HIV-1 JR-FL pseudovirus — NIH AIDS Reagent Program (USA)

DMEM — Gibco™, Thermo Fisher Scientific (USA)

Fetal Bovine Serum — Gibco™, Thermo Fisher Scientific (USA)

Penicillin–Streptomycin — Gibco™, Thermo Fisher Scientific (USA)

Normocin™ — InvivoGen (USA)

Bright-Glo™ Luciferase Assay System — Promega (USA)

VRC01 (bNAb) — NIH AIDS Reagent Program (USA)

#### Criteria for ID_50_ Calculation

ID_50_ values were accepted only when the following predefined criteria were met:

- The serum dilution series spanned the neutralization midpoint, allowing interpolation of 50% inhibition within the tested range [19,27].
- Neutralization data generated an appropriate sigmoidal dose-response curve and demonstrated acceptable goodness-of-fit using a four-parameter logistic regression model [83].
- Positive-control VRC01 neutralization values fell within established assay performance limits [27,84].
- Virus-only and cell-only controls exhibited expected high and baseline RLU values, respectively, confirming acceptable assay performance [27].
- Technical replicates demonstrated acceptable intra-assay variability, and biological replicates showed consistent trends across independent measurements [27,83].

### 4.9. Preclinical Immunogenicity Study

#### 4.9.1. Laying Hens and Production of Anti-HIV-1 gp120 Hyperimmune Egg Yolk

Four healthy laying hens (*Gallus gallus domesticus*), approximately 24–30 weeks of age and actively producing eggs, were used for the generation of anti-HIV-1 gp120 hyperimmune egg yolks. Animals were maintained under standard husbandry conditions with ad libitum access to feed and water throughout the study period. All procedures involving animals were performed in accordance with institutional animal care and ethical guidelines.

The immunogen consisted of commercially available recombinant HIV-1 gp120 envelope glycoprotein corresponding to the HIV-1 IIIB strain (Sino Biological Inc., Beijing, China). For the primary immunization, 100 μg of recombinant gp120 protein was emulsified in an equal volume of Freund’s Complete Adjuvant (FCA) to enhance antigen presentation and antibody production. The emulsion was administered by multiple intramuscular injections into the breast muscles.

Booster immunizations were administered at two-week intervals using 100 μg of recombinant gp120 protein emulsified in Freund’s Incomplete Adjuvant (FIA). A total of three booster immunizations were administered over a six-week period. Following each immunization, hens were monitored for local or systemic adverse reactions, including swelling at the injection site, reduced feed intake, lethargy, or changes in egg production. No significant adverse effects were observed.

Blood samples were collected periodically from the wing vein to monitor the development of the humoral immune response. Serum antibody titres against HIV-1 gp120 were determined by indirect ELISA using recombinant gp120-coated microplates. Seroconversion was defined as a sustained increase in specific anti-gp120 antibody titres above pre-immunization baseline values. Hyperimmune status was considered achieved when endpoint antibody titres exceeded 1:1024, as determined by serial two-fold serum dilutions and geometric mean titre calculations.

Eggs were collected daily beginning two weeks after the first booster immunization and continued throughout the period of peak antibody production. Egg yolks were separated from egg whites, washed extensively to remove residual albumen, and stored at 4°C until processing. Hyperimmune IgY antibodies were subsequently extracted from pooled yolks using the water dilution method described by Polson et al. and were purified for use in the oral immunization experiments.

The concentration and purity of extracted IgY preparations were assessed by spectrophotometric analysis and sodium dodecyl sulfate-polyacrylamide gel electrophoresis (SDS-PAGE). The presence of characteristic IgY heavy chains (approximately 67–70 kDa) and light chains (approximately 25 kDa) was confirmed by electrophoretic analysis. IgY antibodies were isolated from egg yolks according to the procedure described by Polson et al. Briefly, egg yolks were diluted in distilled water under acidic conditions to precipitate lipoproteins, and the water-soluble fraction containing IgY was subsequently separated by centrifugation. The recovered antibody fraction was quantified spectrophotometrically and stored at −20 °C until use [39, 81,82].

#### 4.9.2. Experimental animal: preclinical immunogenicity study

A controlled preclinical immunogenicity study was subsequently conducted. Forty-two healthy outbred male domestic cats (*Felis catus*), aged between 2 and 3 years and weighing approximately 3.0–5.5 kg, were enrolled in this proof-of-concept preclinical study. Animals were obtained from a closed colony maintained under veterinary supervision and were selected to represent the genetic diversity typically observed in domestic feline populations. The use of outbred animals was considered advantageous because it more closely reflects the heterogeneous immune responses expected in naturally occurring populations and may provide a more realistic assessment of vaccine-induced immunogenicity than studies conducted exclusively in genetically homogeneous laboratory animals.

Prior to enrolment, all animals underwent comprehensive veterinary examination, including physical assessment, evaluation of body condition score, temperature measurement, and routine hematological and biochemical screening when available. Animals exhibiting signs of systemic illness, chronic disease, malnutrition, parasitic infestation, oral lesions, respiratory infection, or any other condition potentially affecting immune function were excluded from participation. Only clinically healthy animals were included in the study.

Cats were housed in accordance with institutional animal welfare guidelines in environmentally controlled facilities with adequate ventilation, temperature regulation, and a natural light-dark cycle. Animals had unrestricted access to potable water and were maintained on a commercially formulated feline maintenance diet throughout the study period. Environmental enrichment, including social interaction, toys, and opportunities for physical activity, was provided whenever appropriate to promote animal welfare and minimize stress-related immunological variability.

The study population was divided into two experimental groups. The immunized group (n = 18) received hyperimmune egg yolk preparations containing anti-HIV-1 gp120 IgY antibodies mixed with soy milk, whereas the control group (n = 24) received non-hyperimmune egg yolk preparations mixed with soy milk under identical conditions. Group allocation was performed prior to the initiation of the feeding protocol.

The oral immunization period lasted eight consecutive weeks. Animals were observed daily for evidence of adverse reactions, alterations in feeding behavior, changes in body weight, gastrointestinal disturbances, or other clinical abnormalities. Particular attention was given to signs potentially associated with oral administration of egg-derived immunoglobulins, including vomiting, diarrhea, hypersensitivity reactions, or reduced food consumption. No severe adverse events requiring withdrawal from the study were observed during the experimental period.

Because this investigation was designed as an exploratory proof-of-concept study intended to evaluate the feasibility, safety, and immunogenicity of an oral anti-idiotypic immunization strategy, a formal power calculation was not performed. Instead, the sample size was selected to provide preliminary evidence regarding the induction of anti-HIV-1 immune responses and to generate data for future larger-scale studies.

All experimental procedures involving animals were conducted in accordance with internationally accepted principles for the ethical use of animals in research and were approved by the relevant institutional ethics committees. Animal handling, sample collection, and experimental procedures were performed by trained personnel under veterinary supervision to minimize discomfort and stress. The study adhered to applicable institutional guidelines and internationally recognized standards for animal welfare and biomedical research.

At the completion of the immunization period, saliva samples were collected and analyzed for the presence of anti-HIV-1 gp120 IgA antibodies using an indirect ELISA. The objective of the study was to determine whether oral administration of hyperimmune anti-gp120 IgY could induce antigen-specific mucosal immune responses in outbred felines, consistent with the principles of the common mucosal immune system and mucosal antibody induction following oral antigen exposure [32,33].

### 4.10 Qualitative ELISA for the Detection of Feline Anti-HIV-1 gp120 IgA Antibodies

A qualitative indirect ELISA was developed to detect feline IgA antibodies specific for HIV-1 gp120. Ninety-six-well microplates were coated with commercially available recombinant HIV-1 gp120 protein and blocked with a protein-based blocking solution to minimize non-specific binding. Feline samples (serum, saliva, or other mucosal secretions) were added in duplicate and incubated, followed by the addition of an HRP-conjugated anti-feline IgA antibody. After appropriate washing steps, the enzymatic reaction was developed using tetramethylbenzidine (TMB) substrate and stopped with sulfuric acid. Optical density was measured at 450 nm using a microplate reader.

Positivity was determined using a cut-off value calculated as the mean optical density of negative controls plus two or three standard deviations. Samples with absorbance values above the cut-off were considered positive for anti-gp120 IgA antibodies. This assay allows the qualitative assessment of anti-HIV-1 gp120 IgA antibodies in feline samples and may be useful for studies of mucosal and systemic immune responses, as well as for evaluating immune responses induced by experimental immunization strategies targeting HIV-1 antigens.

### 4.11. Detection of Anti-gp120 IgA Antibodies in Feline Saliva

The induction of mucosal immune responses following oral administration of hyperimmune anti-gp120 IgY was evaluated in a controlled feline immunization study involving 42 outbred domestic cats. Twenty-four animals served as non-immunized controls, while eighteen animals received the oral immunization protocol. All animals were clinically healthy adults aged 2–3 years and were maintained under standard husbandry conditions with unrestricted access to food and water. Animals were monitored daily throughout the study for general health status, behavior, feed intake, and potential adverse reactions associated with oral immunization. Saliva samples were collected at the conclusion of the oral immunization period and analyzed for the presence of anti-HIV-1 gp120 IgA antibodies using a qualitative indirect enzyme-linked immunosorbent assay (ELISA). Ninety-six-well microplates were coated with commercially available recombinant HIV gp120 protein (Sino Biological, Inc), and blocked to minimize nonspecific binding. Saliva samples were tested in duplicate and incubated with horseradish peroxidase (HRP)-conjugated anti-feline IgA antibodies. Following substrate development with tetramethylbenzidine (TMB), absorbance values were measured at 450 nm using a microplate reader. The assay cut-off value was established as the mean optical density of negative control samples plus three standard deviations. Saliva samples were collected at the conclusion of the oral immunization period and analyzed for the presence of anti-HIV-1 gp120 IgA antibodies using a qualitative indirect enzyme-linked immunosorbent assay (ELISA). The detection of antigen-specific salivary IgA was used as an indicator of mucosal immune activation following oral immunization, consistent with current concepts of mucosal immune induction and mucosa-associated lymphoid tissue responses [85].

### 4.12. Ethical Approval

All animal procedures were conducted in accordance with institutional and international guidelines governing the ethical use of animals in research. The study was reviewed and approved by the University of the West Indies, Mona Campus, Jamaica, and by the Higher Institute of Medical Sciences of Santiago de Cuba, Cuba. Documentation of ethical approval was obtained at the time the studies were conducted. All efforts were made to minimize animal discomfort, and animals were maintained under appropriate veterinary care throughout the study [86]. The Campus Ethical Committee of the University of the West Indies at St. Augustine issued the approval numbers CREC-SA.3404/07/2025 and CREC-SA.3430/07/2025.

### 4.13. Statistical Analysis

Data were analyzed using Epi Info™ version 3.5.3 (Centers for Disease Control and Prevention, Atlanta, GA, USA). Descriptive statistics were calculated and are presented as means ± standard deviations (SD), percentages, and geometric mean titers (GMTs) where appropriate. Antibody titers were calculated using the method of Perkins (1958) [87].

For the qualitative anti-gp120 IgA study, differences in the frequency of positive responses between immunized and non-immunized animals were evaluated using Fisher’s exact test [88] because of the categorical nature of the data and the presence of cells with expected counts below five. Odds ratios (ORs) and 95% confidence intervals (95% CIs) were calculated to estimate the strength of association between oral immunization and IgA positivity.

For ELISA optical density measurements, competitive inhibition assays, and HIV-1 neutralization experiments, group comparisons were performed using the Mann–Whitney U test [89] because of the small sample size and the absence of assumptions regarding normal data distribution. Exact p-values were reported whenever possible.

Because this investigation was designed primarily as a pilot proof-of-concept and preclinical immunogenicity study, statistical analyses were intended to identify biologically meaningful trends, estimate effect sizes, and generate preliminary data for future adequately powered studies rather than establish definitive efficacy estimates. Statistical significance was defined as p < 0.05. No adjustments for multiple comparisons were performed because the analyses were exploratory and hypothesis-generating in nature, with the primary objective of identifying biologically meaningful trends and generating preliminary data for future confirmatory studies [**90**].

Exact binomial (Clopper–Pearson) confidence intervals were calculated for proportions where appropriate. Effect sizes including odds ratios and absolute risk differences were calculated to facilitate biological interpretation of the findings. No adjustments for multiple comparisons were performed because the analyses were exploratory and hypothesis-generating.This means that the confidence intervals show precision, the risk difference shows biological magnitude, and the corrected odds ratio shows the strength of association [88,91,92].

To minimize bias, the study incorporated multiple methodological safeguards. Experimental procedures were standardized to reduce variability, including consistent viral input, incubation conditions, among others, which aligns with best-practice recommendations for assay reproducibility [93]. No adjustments for multiple comparisons were performed because the study was designed as an exploratory, hypothesis-generating investigation rather than a definitive confirmatory study.

Randomization of sample placement and blinding of analysts during luciferase quantification helped prevent observer and allocation bias [94,95]. Triplicate measurements and predefined statistical thresholds further reduced analytical bias and improved assay reproducibility [96,97]. The inclusion of appropriate negative, positive, and reference antibody controls ensured internal validity and facilitated the identification of procedural deviations and assay drift [98,99]. Together, these measures strengthened the reliability and interpretability of the experiments.

## 5. Conclusions

The present investigation, integrating a proof-of-concept anti-idiotypic study with a larger preclinical mucosal immunogenicity study, suggests that oral administration of hyperimmune anti-HIV-1 gp120 IgY can induce antigen-specific immune responses in outbred felines. In the proof-of-concept cohort, immunized animals developed detectable anti-anti-idiotypic (Ab3) antibodies, demonstrated modest but specific competitive inhibition, and exhibited measurable HIV-1 neutralizing activity in the TZM-bl assay. Although these findings were obtained in a limited number of animals, they provide preliminary evidence that orally administered anti-gp120 IgY may engage immune network mechanisms capable of stimulating endogenous antibody responses. Furthermore, the observed reduction in luciferase activity in sera from immunized cats was consistent with partial in vitro HIV-1 neutralization under the experimental conditions employed. Collectively, these results support the biological plausibility of anti-idiotypic oral immunization as a strategy for inducing systemic and mucosal immune responses and warrant further investigation in larger studies designed to confirm efficacy, define mechanisms of action, and evaluate translational potential.

A notable finding of this study was the induction of mucosal immune responses in the larger preclinical cohort of 42 outbred cats. Anti-gp120 IgA antibodies were detected in all 18 immunized animals, whereas no positive responses were observed among the 24 control animals. Although these findings should be interpreted within the context of the study design, they suggest an association between oral administration of hyperimmune IgY and the development of antigen-specific mucosal antibody responses. The results are consistent with the possibility that anti-idiotypic oral immunization may influence both systemic and mucosal immune compartments, potentially through pathways involving gut-associated lymphoid tissue and the common mucosal immune system. Further studies will be required to confirm these observations and to elucidate the underlying immunological mechanisms.

Although exploratory in nature and limited by the small size of the proof-of-concept cohort for certain endpoints, the study provides preliminary observations obtained in genetically diverse outbred animals. Collectively, the findings suggest that orally administered hyperimmune IgY may influence immune responses through mechanisms consistent with anti-idiotypic network activation. The detection of mucosal anti-gp120 IgA antibodies, together with the generation of Ab3 antibodies and measurable HIV-1 neutralizing activity in vitro, supports the biological plausibility of this approach and provides a basis for further investigation. Future studies involving larger cohorts, cellular immune analyses, cytokine profiling, epitope characterization, and expanded HIV-1 neutralization testing will be necessary to confirm these findings, clarify the underlying mechanisms, and assess the broader applicability of this immunological strategy.

## Author Contributions

This manuscript was conceptualized by A.J.V. and planning, and research were conducted by all authors. A.J.V and B.F.C. participated in writing the initial draft of the manuscript. Editing and Review: All authors participated. Both authors have read and agreed to the published version of the manuscript.

## Institutional Review Board Statement

The study was conducted under institutional animal-care guidelines in effect at the participating institutions.The Campus Ethical Committee of the University of the West Indies at St. Augustine issued the approval numbers CREC-SA.3404/07/2025 and CREC-SA.3430/07/2025. Instituto Superior de Ciencias Medicas de Santiago de Cuba also approved the use of experimental animals, No. 87-2020.

## Informed Consent Statement

Not applicable.

## Data Availability Statement

The dataset supporting the findings of this study is included in the manuscript and its referenced sources, ensuring comprehensive access to the relevant data for further examination and analysis.

## Disclaimer/Publisher’s Note

The statements, opinions and data contained in all publications are solely those of the individual author(s) and contributor(s) and not of MDPI and/or the editor(s). MDPI and/or the editor(s) disclaim responsibility for any injury to people or property resulting from any ideas, methods, instructions or products referred to in the content.

